# RiboReport - Benchmarking tools for ribosome profiling-based identification of open reading frames in bacteria

**DOI:** 10.1101/2021.06.08.447495

**Authors:** Rick Gelhausen, Teresa Müller, Sarah L. Svensson, Omer Alkhnbashi, Cynthia M. Sharma, Florian Eggenhofer, Rolf Backofen

## Abstract

Small proteins, those encoded by open reading frames, with less than or equal to 50 codons, are emerging as an important class of cellular macromolecules in all kingdoms of life. However, they are recalcitrant to detection by proteomics or *in silico* methods. Ribosome profiling (Ribo-seq) has revealed widespread translation of sORFs in diverse species, and this has driven the development of ORF detection tools using Ribo-seq read signals. However, only a handful of tools have been designed for bacterial data, and have not yet been systematically compared. Here, we have performed a comprehensive benchmark of ORF prediction tools which handle bacterial Ribo-seq data. For this, we created a novel Ribo-seq dataset for *E. coli*, and based on this plus three publicly available datasets for different bacteria, we created a benchmark set by manual labeling of translated ORFs using their Ribo-seq expression profile. This was then used to investigate the predictive performance of four Ribo-seq-based ORF detection tools we found are compatible with bacterial data (Reparation_blast, DeepRibo, Ribo-TISH and SPECtre). The tool IRSOM was also included as a comparison for tools using coding potential and RNA-seq coverage only. DeepRibo and Reparation_blast robustly predicted translated ORFs, including sORFs, with no significant difference for those inside or outside of operons. However, none of the tools was able to predict a set of recently identified, novel, experimentally-verified sORFs with high sensitivity. Overall, we find there is potential for improving the performance, applicability, usability, and reproducibility of prokaryotic ORF prediction tools that use Ribo-Seq as input.

**Key points:**

- Created a benchmark set for Ribo-seq based ORF prediction in bacteria
- DeepRibo the first choice for bacterial ORF prediction tasks
- Tool performance is comparable between operon vs single gene regions
- Identification of novel sORF with DeepRibo is, with restrictions, possible, by using the top 100 novel sORFs sorted by rank.
- Experimental results show that considering translation initiation site data could boost the detection of novel small ORFs
- Determination of novel sORFs in *E. coli* using a new experimental protocol to enrich for translation initiation site. These data-set shows that still a significant part (here 8 out 24, so 1/3) are not detected dispute sufficient Ribo-seq signal. An additional 7 could be recovered using translation initiation site protocols.
- Tools should embrace the use of replicate data and improve packaging, usability and documentation.

## 1 Introduction

Identification and characterization of the proteome is central to understand the biology of viruses and cellular organisms, including bacteria. While mass spectrometry (MS) has been the classical genome-wide approach for protein discovery, it often requires pre-existing open reading frame (ORF) annotations, can be of limited sensitivity, and is strongly influenced by the biochemistry of each protein species. Small proteins (≤ 50 amino acids, aa) are especially difficult to detect by MS [48]. Moreover, the limited information content of small proteins makes them challenging to predict using *in silico* alignment- and sequence-based tools. Small ORFs are likely under-represented in genome annotations [21,38], despite emerging evidence that they play central roles in diverse physiological processes in bacteria, including those underlying virulence [21, 51].

Translation is the last step in protein biogenesis that utilizes RNA, and the power of RNA-seq technology has led to the development of the ribosome profiling (Ribo-seq) approach [27]. Ribo-seq provides a snapshot of the cellular ‘translatome’, which consists of the transcripts that are being actively translated by ribosomes. Ribo-seq coverage therefore serves as a proxy for protein expression. This snapshot is generated by high throughput sequencing of ribosome-protected mRNA fragments that are generated after halting translation by nuclease digestion of RNA not protected by the ribosome. In parallel, the total transcriptome is also sequenced to help to define untranslated regions (UTRs) and estimate the available mRNA input for translation. ORF boundaries can also be defined since Ribo-seq reads are restricted to coding regions. Specific inhibitors that target initiating ribosomes at the start codon (e.g. harringtonine/lactimidomycin in eukaryotes [47] or retapamulin in bacteria [34, 54]) can be used to map translation initiation sites (TISs), which can reveal ORFs hidden within ORFs and increase confidence in the reading frame. In addition to analysis of annotated ORFs, Ribo-seq can also identify novel ORFs missed in genome annotations and proteomic studies. For example, the large number of apparently non-coding transcripts discovered in bacteria by RNA-seq can be re-investigated for their coding potential [40]. Ribo-seq data from diverse organisms, including bacteria, archaea, yeast, mammalian cells, and viruses, has identified a wealth of previously unappreciated coding potential [3, 26, 28, 37, 47, 56].

Despite its power, challenges arise in the analysis of Ribo-seq data to generate robust ORF predictions for downstream characterization. Initially, measures such as translation efficiency (TE), the ratio of ribosome footprint to total transcriptome coverage, were employed to quantitatively detect coding regions. However, this approach can produce high false positive rates [20]. Various groups have therefore developed computational tools that use Ribo-seq coverage patterns and other sequence features for more robust identification of translated ORFs (Table 3). The available tools for ORF prediction from Ribo-seq coverage can be grouped into supervised approaches, such as RiboHMM [43], ORFrater [14], RIBorf [29], REPARATION [36], and DeepRibo [8], or unsupervised methods, such as Ribo-TISH [59], RiboTaper [4], Rp-Bp [32], and SPECtre [7]. Supervised tools require prior training on Ribo-seq data for known/validated ORFs, while unsupervised methods can be used directly for *de novo* predictions. Approaches designed to evaluate the coding potential of transcripts using RNA-seq transcriptome data only, such as CPAT [52], CPC2 [30], and IRSOM [40], have also been developed. Since these cannot use Ribo-seq-specific features like three-nucleotide periodicity, they rely on, e.g., sequence or RNA-seq coverage features. For example, some tools exploit the three nucleotide periodicity in Ribo-seq coverage characteristic of ORFs that arises from movement of the ribosome one codon at a time.

However, experimental challenges have precluded the use of this metric in bacteria [35], and instead bacterial tools have so far relied on detection of coverage and sequence features using machine learning [8, 36]. Bacterial genomes present unique characteristics that can interfere with computational ORF predictions, including high coding density with overlapping genes, unique translation initiation signals, and leaderless transcripts. So far, most tools have been designed and implemented to analyze Ribo-seq data from eukaryotes and do not contain statements about their taxonomic scope, although DeepRibo [8] and REPARATION [36] were developed for and trained on bacterial data. As these tools have not been benchmarked together on bacterial data, their broad utility in these organisms is unclear. While DeepRibo and REPARATION have been compared previously, this was done with the datasets used to train the default model of DeepRibo [8], which are therefore not suitable as an unbiased benchmark set.

Here, we have identified and compared tools for their utility in discovering ORFs from bacterial Ribo-seq datasets, with a special focus on sORFs. First, we generated a novel benchmark ORF set, comprised of an in-house *E. coli* K-12 dataset and three publicly-available datasets for diverse bacterial species. We then used these to quantify and compare the performance of four Ribo-seq-based and one RNA-seq based ORF prediction tool that we found could hand bacterial data. Moreover we tested how well the tools can identify novel, yet unannotated, however experimentally verified sORFs. Finally, we compared tool applicability, usability, and reproducibility to provide a complete picture of their utility.

## 2 MATERIALS AND METHODS

### Ribosome profiling of *E. coli*

#### Growth of bacteria

The *E. coli* MG1655 wild-type strain was grown and harvested for Ribo-seq essentially as described previously [37]. Cultures were grown to mid-log phase (OD_600_ approx. 0.4) in 200 ml lysogeny broth (LB) at 37°C with shaking at 200 rpm. A sample for total RNA was transferred to RNA stop mix (95% ethanol, 5% buffer-saturated phenol (Roth)) and snap-frozen in liquid N_2_. Bacteria were then treated with 100 *μ*g/ml chloramphenicol (final concentration, Sigma) for 2 min at 37°C, followed by harvest via rapid filtration through a 0.45 *μ*m PES (polyethersulfone) membrane (Millipore) and immediate freezing in liquid N_2_.

#### Cell harvest

Harvested cells were processed for Ribo-seq as described previously [37] with minor modifications. Frozen cells were resuspended in chilled lysis buffer (100 mM NH_4_Cl, 10 mM MgCl_2_, 20 mM Tris-HCl, pH 8, 0.1% NP-40, 0.4% Triton X-100, 1 mM chloramphenicol) supplemented with 50 U DNase I (Thermo Fisher Scientific) and 500 U RNase inhibitor (moloX, Berlin) and lysed in Fastprep Lysing Matrix B (MP Bio) for 15 s at speed 4. Clarified lysates (20 A_260_ units) were digested with 2000 U micrococcal nuclease (New England Biolabs) for 1 h (25°C, shaking at 14,500 rpm). Digests were stopped with EGTA (final concentration, 6 mM), immediately loaded onto 10-55% (w/v) sucrose density gradients freshly prepared in sucrose buffer (100 mM NH_4_Cl, 10 mM MgCl_2_, 5 mM CaCl_2_, 20 mM Tris-HCl, pH 8, 1mM chloramphenicol, 2 mM dithiothreitol), and centrifuged (35,000 rpm, 2.5 h, 4°C) in a Beckman Coulter Optima L-80 XP ultracentrifuge and SW 40 Ti rotor. Gradients were fractionated (Gradient Station *ip*, Biocomp) and the 70S monosome fraction (identified by following fraction A_260_) was immediately frozen in liquid N_2_. RNA was extracted from fractions or cell pellets for total RNA using hot phenol:chloroform:isoamyl alcohol or hot phenol, respectively, as described previously [46,50]. Total RNA was digested with DNase I, depleted of rRNA (RiboZero Bacteria, Illumina) and fragmented (Ambion 10X RNA Fragmentation Reagent) according to the manufacturer’s instructions. Monosome RNA and fragmented total RNA was size-selected (26-34 nt) on gels as described previously [25].

#### Library preparation, sequencing, and data deposition

Libraries were prepared by vertis Biotechnologie AG (Freising, Germany) using a Small RNA protocol without fragmentation, and sequenced on a NextSeq500 instrument (high-output, 75 cycles) at the Core Unit SysMed at the University of Würzburg. The results were deposited in the NCBI Gene Expression Omnibus (GEO) and are available via GSE131514.

### Public data retrieval

#### Escherichia coli

K-12 MG1655: Published proteomics data [45] were obtained from [Supplemental Table S9](https://static-content.springer.com/esm/art%3A10.1038%2Fnbt.3418/MediaObjects/41587_2016_BFnbt3418_MOESM18_ESM.xlsx) of the cited manuscript. Cultures were grown at 37°C in LB until they completed ten divisions in exponential state. In order to test the ability of the tools to detect novel sORFs, we retrieved an additional *E. coli* MG1655 dataset, distinct from our newly generated dataset. We retrieved published [54] Ribo-seq (SAMN10583712, SAMN10583713) dataset for bacteria grown at 37°C in MOPS EZ Rich Defined media with 0.2% glucose to an OD_600_ of 0.3. Experimentally verified novel ORFs were retrieved from Table 1 of the publication.

**Table 1:** Overview of identified ORF detection tools. Most tools make no statement about the taxonomic domain they were developed for, some however utilize eukaryotic organisms as example datasets (indicated by *). The first five tools were benchmarked in this manuscript.

| Name | Input data | Method | Availability | Taxonomy |
| --- | --- | --- | --- | --- |
| DeepRibo [8] | Ribo-seq | Deep Learning | github | Prokaryotes |
| Reparation_blast [36] | Ribo-seq | Random Forest | bioconda, github | Prokaryotes |
| SPECTre [7] | Ribo-seq | Spectral Coherence | github | Eukaryotes* |
| Ribo-TISH [59] | Ribo-seq | Negative Binominal Test | bioconda, github | Eukaryotes* |
| IRSOM [40] | RNA-seq | Self-Organizing Map | gitlab, webservice | Eu-, Prokaryotes |
| RiboTaper [5] | Ribo-/ RNA-seq | Multitaper Spectral Analysis | bioconda, galaxy | Eukaryotes* |
| RiboHMM [43] | Ribo-, RNA-seq | Hidden Markov Models | github | Eukaryotes |
| ORFrater [14] | Ribo-seq | Linear Regression | github | Eukaryotes* |
| RibORF [29] | Ribo-seq | Logistic Regression | github | Eukaryotes* |
| Rp-Bp [32] | Ribo-seq | Markov Chain - Monte Carlo | github | Eukaryotes* |

#### *Listeria monocytogenes* EDG-e

For *L. monocytogenes*, we utilized data from a published screen for antibiotic-responsive ribo-regulators [10]. We retrieved the Ribo-seq (SAMEA3864955) and RNA-seq (SAMEA3864956) datasets for the wild-type strain EDG-e from SRA. Cells were grown in brain heart infusion (BHI) medium at 37°C to an OD_600_ of 0.4-0.5. The culture was supplemented with control medium for 15 min before harvesting. For our analysis, the untreated control library was used. Published proteomics data [24] was obtained from Supplemental Tables [S2,3,4,5,6,7,8](https://www.ncbi.nlm.nih.gov/pmc/articles/PMC5802382/bin/NIHMS75486-supplement-Supplementary_table_S2.xlsx) of the cited manuscript. Cultures were grown at 37°C to an OD_600_ of 1.

#### *Pseudomonas aeruginosa* PAO1

The data for *P. aeruginosa* is from a study investigating expression differences in strains with high sequence similarity but differences in substrate consumption efficiency using a multi-omics approach [18]. We retrieved the Ribo-seq and RNA-seq (SAMN06617371) datasets for the PAO1 wild-type strain grown on n-alkanes to mid-log phase. Corresponding proteomics data was retrieved from Supplemental Tables [S21-S24](https://www.ncbi.nlm.nih.gov/pmc/articles/PMC5410065/bin/12864_2017_3708_MOESM6_ESM.xlsx) of the same publication.

#### Salmonella

Typhimurium 14028s: Finally, we used data generated to investigate the impact of the RNA-binding protein CsrA on *S.* Typhimurium virulence-associated stress responses and metabolism [41]. We retrieved Ribo-seq (SRX3456030) and RNA-seq (SRX3456038) datasets for wild-type strain 14028s grown in LB medium at 37°C to an OD_600_ of 0.5. The published [58] MS data was obtained from Supplemental Table [S1](https://www.ncbi.nlm.nih.gov/pmc/articles/PMC3213010/bin/1752-0509-5-100-S1.XLSX) of the cited manuscript. Cultures were cultivated under identical conditions as for Ribo-seq.

### Bioinformatic analysis

In order to evaluate the ORF detection tools, we used part of a pre-release version of our HRIBO (High-throughput annotation by Ribo-seq) workflow [16], which we have developed to analyse prokaryotic ribosome profiling experiments [51]. The genomes and annotations of *E. coli* K-12 substr. MG1655 (ASM584v2), *L. monocytogenes* EGD-e (ASM19603v1), *P. aeruginosa* PAO1 (ASM676v1, ASM75657v1), and *S.* Typhimurium 14028s (ASM2216v1) retrieved from the National Center for Biotechnology Information (NCBI) [12] were used. The HRIBO workflow consists of 3 steps: the pre-processing of the input data, the execution of the individual prediction tools, and a post-processing step. A detailed description of how to run the RiboReport pipeline is provided in the [RiboReport](https://github.com/RickGelhausen/RiboReport) GitHub repository. The individual steps of the workflow are described in the following.

#### Pre-processing

To generate the required input files for the benchmarking tools, adaptors were first trimmed from the input reads using cutadapt [33]. Next, reads were mapped to the genome using segemehl [22], which has higher sensitivity than other mappers, and its high computational costs are still acceptable for small genomes. Finally, the reads mapping to ribosomal RNA or multiple genomic locations were filtered out using samtools [31]. Adapted annotation files were also generated, as several tools require very specific formatting of *GTF* files. DeepRibo, requires coverage files as an input. The coverage files were produced using a custom-made script, following the instructions in the DeepRibo documentation [8]. In summary, we generated read alignments to the respective reference genomes for Ribo-seq and RNA-seq libraries in *BAM* (Binary version of sequence alignment map format), as well as transcript files in *BED* (Browser Extensible Data) and read coverage files in *BEDGRAPH* format. In addition, we monitored the quality of each of these steps using fastQC and aggregated the results into a MultiQC [13] report.

#### Execution of ORF detection tools

Tools compatible with bacterial data and annotations were investigated: Ribo-TISH, Reparation_blast, DeepRibo, SPECtre, and IRSOM. Since all tools, with the exception of Ribo-TISH, do not handle replicates, we selected a single replicate for each organism. Ribo-TISH was called using default parameters using the mapping files generated from the Ribo-seq data, the reference genome, and the adapted annotation file. Since Ribo-TISH could not use reference annotations from NCBI [12], we generated an annotation file that only contains *gene*, *exon*, and *CDS* features and *gene_id* and *transcript_id* attributes. Reparation_blast was run using default parameters with the Ribo-seq mapping files, the reference genome and annotation, and the uniprot_sprot [9] database. Since REPARATION uses the commercial tool ublast internally, we replaced ublast with *protein blast* (blastp) [6] and adapted the tool to allow the input of *BAM* files. Since blastx is more sensitive while consuming more CPU-time compared to ublast [57], we expect that our modified tool behaves similarly in comparison to the original version. We made this adapted version, called Reparation_blast, available via *bioconda* [19]. SPECtre was executed with default parameters, using a isoforms file created by cufflinks. SPECtre accepts annotation files only as ensembl-formated general feature format (v2, *GTF*) files, which made it necessary to convert the NCBI general feature format (v3, *GFF*) files used for the other tools using a custom script.

DeepRibo parameters for noise reduction need to be adapted for each data set. We used the script provided by the DeepRibo GitHub repository (*s_curve_cutoff_estimation.R*) for this purpose. This script provides cut-off values for *coverage* and RPKM (reads per kilobase million). Furthermore, we provided it with the requested input coverage and acceptor site coverage files, as well as the reference annotation, the reference genome, and the included pre-trained model. IRSOM was called using default parameters and the included pre-trained model for *E. coli*. All other pre-trained models are dedicated to the use of eukaryotic organisms. Further, we used cufflinks [49] to extract transcript regions from the alignment files generated from RNA-Seq data and provide these to IRSOM for prediction.

#### Post-processing

Post-processing steps were performed by parsing the prediction results of each tool into a *GTF* format file that can be used for evaluation. As each tool has a different output format, each result file had to be parsed differently. For Reparation_blast and SPECtre, we converted the results from a text file into *GTF* format. For Ribo-TISH we used the *RiboPStatus* column to select only the best result for each start codon. For DeepRibo we used the *SS_pred_rank* column to select only the best result for each stop site. Finally, for IRSOM, which reports whether a result is coding or non-coding, we only used results labeled as coding. Additionally, the workflow generates multiple excel files containing different measures, like translational efficiency, RPKM, amino acid count, and others. These files were used in order to assist with the manually labeled dataset of the annotated features.

#### Processing of mass spectrometry data

Mass spectrometry data was first converted to *GFF* format. The exact steps required for the different datasets can be reproduced as described in the RiboReport [proteomics](https://github.com/RickGelhausen/RiboReport/tree/master/proteomics) directory.

### Benchmark of ORF detection

#### Labeling of translated regions based on Ribo-seq data

We tested the predictive power of the tools using annotated genes within the NCBI annotation. For each organism, a human expert (S.L.S.) made judgments about whether each annotated ORF is translated/not-translated as follows. One RNA-seq replicate and its corresponding Ribo-seq (70S footprint) library were loaded into the Integrated Genome Browser (IGB) [15] together with the genome reference sequence and ORF annotation. Each ORF feature was visually inspected without knowledge of the locus tag or gene product name. We labeled ORFs as translated by using following criteria. The coverage of loci in RNA-seq libraries and Ribo-seq libraries was inspected. ORFs were required to have a coverage in the Ribo-seq library of at least 10 RPKM. The Ribo-seq signal needed to be at least as strong as the signal of the transcriptome library (i.e., TE >1). Moreover, the ORFs needed to be enriched near the start codon and/or within the ORF boundaries. For manual labeling of the 33 western blot validated sORFs from [54], the same approach was taken, with the exception that only the Ribo-seq library was inspected as no RNA-seq library was provided with the dataset. The associated TIS library is only included in screenshots, and was not used for the manual labeling.

#### Computation of prediction quality

For each organism, we used the manually labeled datasets (*labels.gff*) to split the ORFs into two files (*positive_labels.gff*, *negative_labels.gff*) representing translated and non-translated ORFs, respectively. The set of condition-positive ORFs (those labeled as translated in our manual curation) should therefore be found by a prediction tool, while the condition-negative ORFs (those labeled as not translated) and should not be called as translated).

To determine whether a prediction should be assigned to an annotated ORF from our benchmark set, we defined different overlap thresholds between the genomic coordinates of a prediction and the ORFs labeled as translated or non-translated. The overlap was computed using bedtools intersect [42].

We set reciprocal overlap thresholds of 1%, 70% and 90%, requiring that the label-prediction overlap, and vice versa, is at least as big as the selected threshold. For example, the overlap threshold of 1% tests whether a tool detects translation at a certain locus at all, while the 90% threshold tests if a tool can also predict its correct length. We decided to use a threshold of 70% to emulate the inspection strategy of a researcher who will inspect ORFs of interest afterwards. This cutoff tests for translation of a locus, but includes the possibility to identify novel truncated or nested ORFs.

Based on the intersection between the tool predictions and our manually labeled ORF sets, each ORF prediction was classified as a true positive (TP), true negative (TN), false positive (FP), or false negative (FN). An annotated gene with a positive label was counted as a TP if there was at least one prediction that was associated with the gene, and as a FN if no prediction was associated with the gene. An annotated gene with a negative label was counted as an FP if there was at least one prediction associated with the gene or a TN if no prediction was associated with the gene. The association of predictions and genes was determined for each tool and dataset individually. There were two cases where a prediction was not counted for a labeled gene. First, an annotated gene might have an overlap with multiple predictions from a given tool. In this case, only the prediction with the best predictive score or probability, depending on the tool, was considered. All other predictions were counted as sub-optimals and ignored for the remaining analysis. Second, there were predictions that did not overlap with any annotated ORFs. These predictions were not counted at all, as the ground truth is not known in this case (i.e., we cannot determine whether they were novel predictions or FPs).

In addition to comparing the tools for the *E. coli* NCBI ORF annotation, we also investigated their performance on novel sORFs using a Ribo-seq dataset for *E. coli* that was generated in parallel with a TIS library that revealed 33 novel sORFs that were independently validated by western blotting (see subsection *Novel small ORFs*).

To measure the prediction quality of the tools in determining the correct labels for each ORF of our benchmark, we computed the sensitivity and specificity of their predictions. Since our positive and negative datasets were unbalanced, we computed the F1 measure as an unbiased tool performance measurement. Furthermore, we plotted Precision-Recall Curves (PRC) and calculated their area under the curve (AUC) to compare the performance of the different tools between the organism. The PRC avoids an overlap threshold bias, unlike the F1 measure, which can only be calculated for one overlap threshold. To compute PRCs, the positively- and negatively-labeled ORFs were used to generate the positive and negative datasets, respectively. Since the computed scores of the tools were not directly comparable, all predictions were ranked based on their given scores. Annotated ORFs without an associated prediction (FN and TN) were included in the ranking with the lowest possible score that each tool could provide.

Evaluation scripts are located in the evaluation directory of the [RiboReport](https://github.com/RickGelhausen/RiboReport/tree/master/evaluation) repository, with a [description](https://github.com/RickGelhausen/RiboReport#visualisation-and-plotting-of-the-results) on how they were executed. The PRC and AUC were computed using scikit-learn [23] and plotted using matplotlib [39]. In addition to the PRC, each plot includes a baseline (baseline = positive labels/(positive labels + negative labels)), which represents how many positive predictions are expected to occur by chance. For each Venn diagram, overlap sets of the correctly-discovered, positively labeled ORFs were computed. We used the Jvenn webserver to produce the Venn diagrams [2] in Figures 2, 3 and python scripts utilizing the seaborn [53] and simple_venn library for Figure S1.

#### Selection of subsets

Besides the whole *translatome* dataset, we also tested tool performance on the following subsets: (1) *Operons* were defined as groups or intervals of neighboring genes on the same strand with an intergenic distance of less than 200 nucleotides (see here for [description](https://github.com/RickGelhausen/RiboReport#4-extract-operon-regions-from-the-annotation)). (2) *Non-operon ORFs* are those that do not overlap with the *Operons* intervals. (3) *Small ORFs* were defined as genes with length ≤ 150*nt* (50 aa) [54]. Based on these definitions, we generated labeled positive (translated) and negative (not translated) sets for each subset. These files are available in our [GitHub repository](https://github.com/RickGelhausen/RiboReport/tree/master/data).

#### Computation of run time and peak memory consumption

Runtime and memory consumption of the tools was evaluated by running the tools individually on our newly generated *E. coli* dataset with either a single or with ten CPU threads. The machine used for testing featured an AMD EPYC 7351P 16-Core processor, 130 GB RAM, and was running Ubuntu 18.04.1 (kernel version 4.15.0-76-generic).

#### Evaluation of manual labeling with mass spectrometry data

To validate our labeling method, each annotated ORF in the four bacterial genomes was first manually labeled as translated or not based on manual inspection of Ribo-seq data in a genome browser (see Methods for details). We then validated our labeling approach by comparison to available published MS datasets ([proteomics](https://github.com/RickGelhausen/RiboReport/tree/master/proteomics)) for the same strains grown under similar conditions (see Supplement Section - Validation of labeling method, Figure S1). The MS data was selected to be as as similar as possible to the Ribo-seq experimental conditions (Data Retrieval).

## 3 RESULTS & DISCUSSION

### Available ORF prediction tools and their applicability to bacterial data

By screening a review [5] and recently published studies [8, 36], we found nine Ribo-seq based ORF detection tools(3). Additionally, we identified several tools that predict potential ORFs from only RNA-seq (transcriptome) data and included the newest example (IRSOM) for comparison. We first tested the ten identified ORF detection tools for their compatibility with bacterial annotations using our *E. coli* benchmark dataset. We found that only five tools could accept and process these datasets: Reparation_blast, Ribo-TISH, IRSOM, SPECtre, and DeepRibo. Since RiboTaper and RiboHMM do not work with bacterial annotations, we could not run them. We were not able to install Rp-Bp on our cluster system or locally in a reasonable amount of time. For ORFrater and RIBorf, several steps of their pipelines could be executed, but we did not obtain a result output.

The five tools that could be used with bacterial data take different approaches to ORF detection. DeepRibo [8] was designed for bacteria and uses a convolutional network with a one-hot encoding of the DNA sequence to detect important sequence motifs such as the Shine-Dalgarno sequence. In this case, the one-hot encoding converts the sequence into a binary representation. This network is then combined with a recurrent neural network architecture to model the periodicity found in the Ribo-seq data. DeepRibo models have been trained on multiple Ribo-seq experiments from different bacterial species. REPARATION [36] was also developed for bacterial data and trains a random forest classifier on all possible ATG-, GTG- and TTG-starting ORFs. Candidates below a minimum RPKM cutoff or footprint coverage are considered as noise and removed from the prediction. This noise level is determined from the lower bend point of a sigmoid curve fitted to the RPKM/coverage plot. The classifier is then used on all potential ORFs satisfying the thresholds. REPARATION uses the proprietary homology search tool ublast [11] internally, which we replaced by the open tool blastp [6] to make REPARATION viable for open source usage, e.g. in pipelines. We refer to this version as Reparation_blast.

Two of the tools are designed for eukaryotic Ribo-seq data. Ribo-TISH [59], developed for eukaryotes, tests reading frames with a non-parametric Wilcoxon rank-sum test on the read count difference for each ORF nucleotide position. The test is used to distinguish between the alternative reading frames to determine the translated ORF. SPECtre [7] is based on spectral coherence to predict regions of active translation from mapped Ribo-seq data. It matches the periodic reading frame function with the signal of aligned reads, using a Welch’s spectral density estimate to compute SPECtre scores. Distributions of these scores are then used to assign a posterior probability predicting if a given region is translated.

In addition to the tools that accept/processed on Ribo-seq data, we tested the performance of IRSOM [40], which predicts coding regions from RNA-seq data and was established in eukaryotes. IRSOM uses multiple features extracted from the candidate, such as read distribution over different regions of the ORF, as well as length and reading frame properties. Additionally, sequence features, e.g. nucleotide and k-mer motif frequencies, GC content, and codon properties, are used to create a supervised classifier based on self-organizing maps with a fully connected perceptron layer. IRSOM was included in the benchmark to investigate the additional predictive power gained by using Ribo-seq data.

### Benchmark datasets

A robust performance evaluation of sORF detection tools requires data from a variety of prokaryotic organisms. Therefore, we selected multiple publicly-available datasets covering different bacterial species for our benchmark. Criteria for selection included quality (published, sufficient sequencing quality, sufficient documentation (*i.e.*, adaptor sequences)), as well as the availability of a paired RNA-seq library to aid manual labeling of ORF translation and for evaluation of RNA-seq-based IRSOM. In addition, we added our own *de novo*-generated *E. coli* dataset. In total, the four benchmark datasets include our newly-generated *E. coli* dataset and three publicly available datasets for wild-type strains of *L. monocytogenes*, *P. aeruginosa*, and *S.* Typhimurium (Table 2) (see Methods for details). These manually labeled Ribo-seq ORF sets are, to our knowledge, the first available for bacterial Ribo-seq data for the purpose of tool benchmarking. The manually labeled Benchmark sets for the four organisms are available from the [GitHub repository](https://github.com/RickGelhausen/RiboReport/tree/master/data). Description of labeling quality via comparison to MS data and inspection of specific examples is described in the Supplemental Materials (see Supplemental Figs. S1-S3).

**Table 2:** Generation of a curated benchmark ORF set. The benchmark set archives contains *GFF* files for labels of all annotated ORF sets (positive/negative), MS labels, tool predictions, operons, genome sequence and reference annotation to enable inspection in the genome browser. Links to the original data sources are provided. The number of ORFs from each annotated ORF set (*translatome*, *small ORFs*, *operon*, and *non-operon*) that have been identified as translated (positive) or non-translated (negative) are listed.

| Organism | *E. coli* |  | *L. monocytogenes* [10] |  | *P. aeruginosa* [41] |  | *S. Typhimurium* [18] |  |
| --- | --- | --- | --- | --- | --- | --- | --- | --- |
| Benchmark set [zip] | E. coli |  | L. monocytogenes |  | P. aeruginosa |  | S. Typhimurium |  |
| Growth conditions | WT, LB @ 37°C |  | WT, BHI @ 37°C |  | WT, n-alkanes |  | WT, LB @ 37°C |  |
| Data | GSE131514 |  | SAMEA3864955<br>SAMEA3864956 |  | SAMN06617371 |  | SRX3456030<br>SRX3456038 |  |
| Set | positive | negative | positive | negative | positive | negative | positive | negative |
| Translatome | 2765 (65%) | 1483 (35%) | 2288 (80%) | 579 (20%) | 3938 (71%) | 1635 (29%) | 3290 (66%) | 1681 (34%) |
| Small ORFs | 55 (48%) | 59 (52%) | 7 (100%) | 0 (0%) | 7 (58%) | 5 (42%) | 31 (31%) | 69 (69%) |
| Operons | 1795 (64%) | 1014 (36%) | 1622 (80%) | 432 (20%) | 2512 (69%) | 1112 (31%) | 1953 (66%) | 1002 (34%) |
| Non-operons | 970 (67%) | 469 (33%) | 666 (82%) | 147 (18%) | 1426 (73%) | 523 (27%) | 1337 (66%) | 679 (34%) |

### Benchmark results

DeepRibo and Reparation_blast have been recently compared for their performance [8]. However, this comparison was based on a dataset used to train the default model of *DeepRibo*, this is therefore not an unbiased benchmark. We thus used our novel, comprehensive benchmark set to evaluate the performance of all five ORF detection tools that we found accept bacterial data (Table 3). Prediction quality metrics were computed (see Methods subsection Benchmark of ORF detection) for the whole *translatome*, as well as for specific ORF subsets that have properties that could possibly influence prediction results. We compared whether the predictors show a different behaviour inside and outside of operon regions, as well as for annotated sORFs and a set of western blot validated novel sORFs from *E. coli* using an additional Ribo-seq dataset [54].

**Table 3:** Overall tool performance at different overlap thresholds. The AUC of the precision-recall curve is given for the tools with each of the four organism datasets (whole *translatome* ORF set) at the prediction overlap thresholds of 1%, 70% and 90%. The overlap threshold is the percentage of the ORF length that the prediction must satisfy.

| Organism | *E. coli*<br>(AUC) |  |  | *L. monocytogenes*<br>(AUC) |  |  | *P. aeruginosa*<br>(AUC) |  |  | *S. Typhimurium*<br>(AUC) |  |  |
| --- | --- | --- | --- | --- | --- | --- | --- | --- | --- | --- | --- | --- |
|  | 1% | 70% | 90% | 1% | 70% | 90% | 1% | 70% | 90% | 1% | 70% | 90% |
| DeepRibo | 0.97 | 0.96 | 0.95 | 0.88 | 0.88 | 0.88 | 0.95 | 0.95 | 0.95 | 0.97 | 0.96 | 0.95 |
| Reparation_blast | 0.82 | 0.82 | 0.82 | 0.94 | 0.94 | 0.94 | 0.9 | 0.9 | 0.89 | 0.94 | 0.96 | 0.96 |
| Ribo-TISH | 0.85 | 0.60 | 0.60 | 0.83 | 0.75 | 0.75 | 0.85 | 0.68 | 0.65 | 0.87 | 0.73 | 0.73 |
| IRSOM | 0.67 | 0.67 | 0.67 | 0.78 | 0.77 | 0.77 | 0.67 | 0.67 | 0.67 | 0.68 | 0.69 | 0.69 |
| SPECTre | 0.65 | 0.65 | 0.65 | - | - | - | 0.67 | 0.67 | 0.67 | 0.71 | 0.71 | 0.71 |

#### Bacterial tools generally show more robust performance

The tools were first compared on the whole complement of annotated ORFs for each organism (hereafter the *translatome* set) (Table 2). Tool performance was measured by determining the area under the curve (AUC) of a Precision-Recall Curve (PRC) [44]. We selected this metric because the number of positively- and negatively-labeled ORFs were imbalanced, especially for *L. monocytogenes* (80% of ORFs were in the positive set). The PRC compares the recall of the tool against its precision value for a given score cutoff. The recall in this context is the fraction of correctly-predicted, labeled ORFs (true positives, TPs) versus the sum of all positively-labeled ORFs (including false negatives (FN)), yielding (TP/TP + FN). The precision is the fraction of correctly-predicted, positively-labeled ORFs (TPs) versus the sum of all positively-predicted ORFs (including FP) yielding (TP/TP + FP). We compared the AUC for each tool at different overlap thresholds to test not only if they were able to predict the presence of an ORF, but also if they could correctly determine its length (Table 3). We used thresholds of 1%, 70%, and 90% (i.e., the prediction must cover at least 1%, 70%, 90% of the ORF length). DeepRibo, Reparation_blast, SPECtre and IRSOM showed a stable performance over the three thresholds, meaning that when they predict an ORF they also can correctly predict its length. Ribo-TISH, however, often predicted only a short region of the annotated ORF as translated. This can be observed, for example, in *E. coli*, where the high AUC of 0.85 for the 1% overlap threshold then drops to an AUC of 0.6 for the 70% overlap threshold. The PRCs for a overlap threshold of 70% (Figure 1) show that DeepRibo and Reparation_blast performed well for detection of the *translatome* benchmark ORF set from all four organisms (AUC < 0.8). In contrast, IRSOM, SPECtre, and Ribo-TISH generally had substantially lower AUCs - almost close to random (grey baseline, see methods subsection: Benchmark of ORF detection).

**Figure 1:**
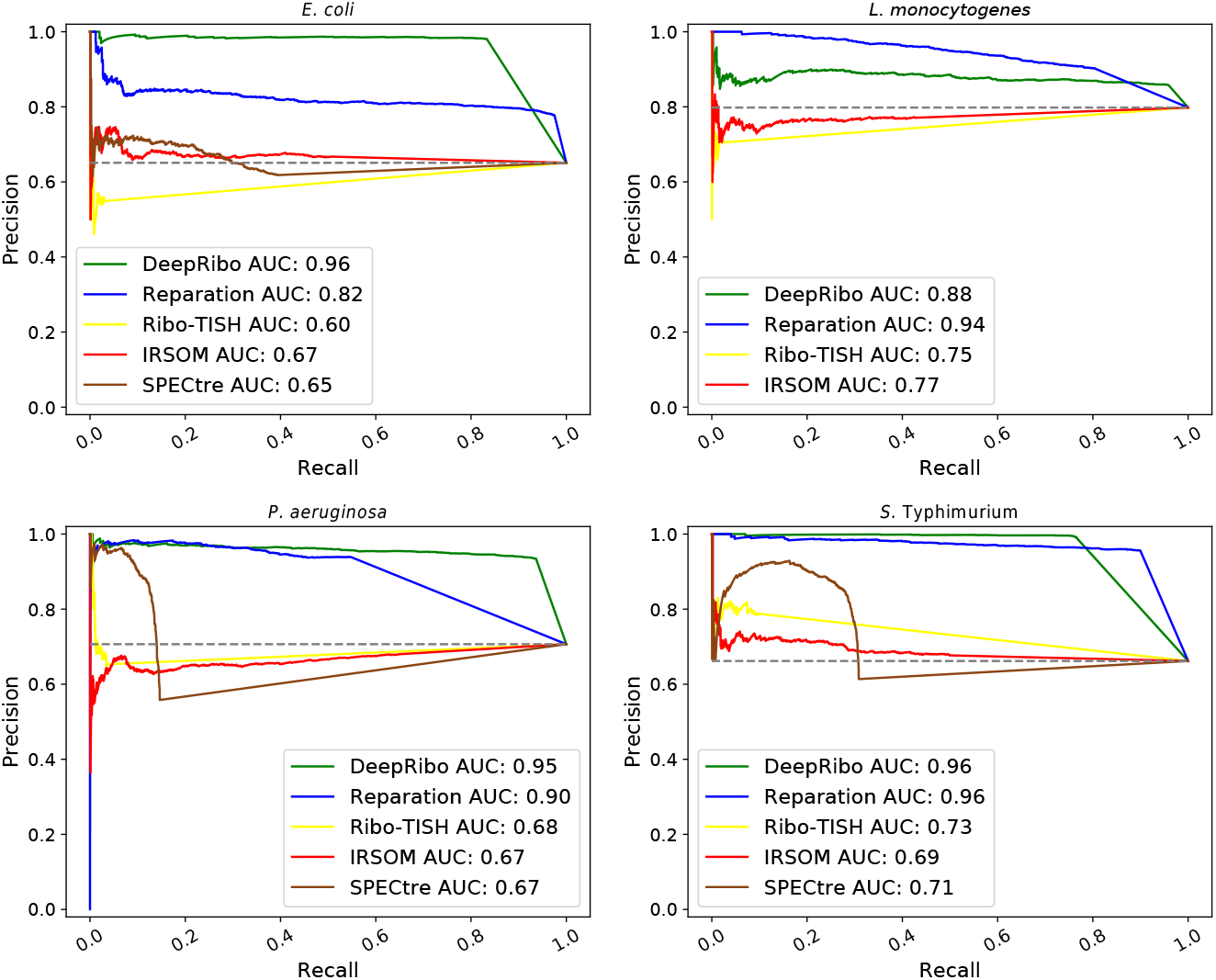
Precision-Recall curves (PRCs) for *E. coli*, *L. monocytogenes*, *P. aeruginosa* and *S.* Typhimurium. Predictions were ranked according to the score provided by each tool (e.g., the probability for Reparation_blast or the prediction rank for DeepRibo). A prediction was associated with a labeled ORF if more than a 70% overlap exists between the sequence of the prediction and the labeled ORF, and vice versa. All associated instances (positive and negative dataset) were ranked according to there scores. The ranked instances were than used to plot the PRC and to calculate the AUC. The grey baseline indicates how many predictions are expected to occur by chance.

DeepRibo showed the highest AUC values for *E. coli*, *S.* Typhimurium, and *P. aeruginosa*, suggesting it has the highest predictive power for most organism datasets, while Reparation_blast performed best for *L. monocytogenes*. A possible explanation for this is that the organisms DeepRibo was trained on might have different genomic characteristics compared to *L. monocytogenes*. However, it could also be the result of experimental differences that change the distribution of the read coverage.

We next investigated the sensitivity, specificity, and F1 measure of the tools (Table 3). The F1 measure, which is the harmonic mean of recall and precision, showed that IRSOM performed surprisingly well, even though it only relies on RNA-seq data. IRSOM, however, could not compete with the tools designed for bacterial Ribo-seq data (DeepRibo and Reparation_blast). This same trend was observed for sensitivity and specificity. DeepRibo showed overall a strong predictive performance, only being outperformed by Reparation_blast for the *L. monocytogenes* data set. The lower AUC value in this case is due to a higher FP rate for this data set (see Supplement Tables 1-4).

The sensitivity of Ribo-TISH was low for all four datasets (Table 3). As already seen for the AUC at different overlap thresholds (Table 3), Ribo-TISH did not predict ORFs precisely, but rather predicted a short signal nested in the region of a labeled ORF (average sensitivity for overlap threshold 0.01 is 0.6). SPECtre, similar to Ribo-TISH, had a low sensitivity. However, its specificity while comparable is slightly lower. We could not obtain SPECtre results for *L. monocytogenes*, since the prediction did reproducibly not terminate within 72h. The lower performance of Ribo-TISH and SPECtre might be explained by the fact that they were not specifically designed for bacteria, which have distinct translatome structures. In addition, both tools rely on three nucleotide periodicity, which is often not pronounced in bacterial datasets due to experimental issues. Moreover, SPECtre depends on the transcript-calling performance of cufflinks [49], which means that it is also affected by the quality of coupled RNA-seq data.

In addition to computation of global performance metrics, we also qualitatively compared how the tools performed for specific ORFs. We inspected coverage for specific examples of ORFs in genomic regions conserved between the four benchmark organisms and compared this to their detection by each of the five tools at a 70% overlap threshold. For this and future comparisons, genome browser tracks for all tool predictions can be found as prediction.gff files in the archives of each respective organism [RiboReport](https://github.com/gelhauser/RiboReport) repository (data/*/misc *.zip, * = organism). We first compared the detection of genes in a ribosomal protein island with conserved synteny to assess our labeling performance vs MS (Supplemental Figure S2), all of which are likely *bona fide*, translated ORFs under the conditions tested due to their central role in translation. Comparison of detection by the five tested tools showed that in general, DeepRibo and Reparation_blast called these ORFs as translated. In comparison, SPECtre and Ribo-TISH did not detect any of the 22 ORFs at this threshold. Surprisingly, RNA-seq based IRSOM was mildly successful, detecting a handful of ORFs in the organisms other than *E. coli*. We also examined tool predictions of two genes in an operon shared by all four bacteria: that encoding a terminal oxidase (*cydAB* in *E. coli*, *S.* Typhimurium, and *L. monocytogenes*, *cioAB* in *P. aeruginosa*) (Supplemental Figure S4). Both *cydA* and *cydB* were labeled as translated and detected by DeepRibo in all organisms, while Reparation_blast detected all but *cioAB* in *P. aeruginosa*. The other tools showed variable detection of the *cydA*/*cydB* homologues, with Ribo-TISH not detecting either in any organism. Closer inspection of the predictions (data not shown) indicated that Ribo-TISH was predicting several very short nested ORFs in *cydA* and *cydB*. Together, these comparisons of tool sensitivity and specificity on the whole translatome ORF sets for each of the four bacterial species shows that the bacterial Ribo-seq tools are superior to IRSOM and Ribo-TISH.

#### Operon-encoded ORFs

A unique feature of bacterial genomes is the operon structure: several genes, often of related function, are transcribed as one polycistronic mRNA. Operons often have small distances between ORFs that might lead to ambiguity in associating Ribo-seq signal between neighboring ORFs. They might even include overlap of coding regions. These features could presumably affect ORF prediction tools. Therefore, we tested whether the predictive power of the tested tools is different for ORFs within operons compared to single transcribed genes (Table 5).

**Table 4:** Detailed tool performance measures for 70% overlap. The sensitivity or true positive rate (TPR), specificity or true negative rate (TNR), and the F1 measure were calculated for each tool with each organism benchmark dataset (*translatome*). The sensitivity highlights how well the positive labels are detected and the specificity reveals how well negatively labeled ORFs are not predicted by the tools. The F1 measure is an unbiased tool accuracy measurement. The values were calculated with the requirement that the prediction of an ORF must be covered by at least 70% of its coding sequence.

| Organism Measure | *E. coli* |  |  | *L. monocytogenes* |  |  | *P. aeruginosa* |  |  | *S. Typhimurium* |  |  |
| --- | --- | --- | --- | --- | --- | --- | --- | --- | --- | --- | --- | --- |
|  | TPR | TNR | F1 | TPR | TNR | F1 | TPR | TNR | F1 | TPR | TNR | F1 |
| DeepRibo | 0.83 | 0.97 | 0.90 | 0.96 | 0.37 | 0.91 | 0.94 | 0.84 | 0.94 | 0.77 | 0.98 | 0.86 |
| Reparation_blast | 0.98 | 0.48 | 0.87 | 0.81 | 0.65 | 0.85 | 0.55 | 0.91 | 0.7 | 0.9 | 0.92 | 0.93 |
| Ribo-TISH | 0.03 | 0.95 | 0.06 | 0.02 | 0.96 | 0.05 | 0.04 | 0.96 | 0.07 | 0.1 | 0.95 | 0.17 |
| IRSOM | 0.51 | 0.53 | 0.58 | 0.43 | 0.5 | 0.55 | 0.66 | 0.25 | 0.67 | 0.5 | 0.53 | 0.58 |
| SPECTre | 0.39 | 0.55 | 0.48 | - | - | - | 0.15 | 0.72 | 0.23 | 0.31 | 0.62 | 0.41 |

**Table 5:** Prediction of ORFs in operons. The predictive power of the five tools for translation of genes either inside of polycistronic regions (operon) or in monocistronic regions (non-operon) was compared via the AUC for precision-recall curves computed using an overlap threshold of 70%.

| Organism ORF type | *E. coli* (AUC) |  | *L. monocytogenes* (AUC) |  | *P. aeruginosa* (AUC) |  | *S. Typhimurium* (AUC) |  |
| --- | --- | --- | --- | --- | --- | --- | --- | --- |
|  | operon | non-operon | operon | non-operon | operon | non-operon | operon | non-operon |
| DeepRibo | 0.96 | 0.96 | 0.86 | 0.91 | 0.95 | 0.95 | 0.96 | 0.96 |
| Reparation_blast | 0.82 | 0.82 | 0.93 | 0.95 | 0.91 | 0.89 | 0.96 | 0.96 |
| Ribo-TISH | 0.59 | 0.62 | 0.74 | 0.77 | 0.65 | 0.73 | 0.71 | 0.76 |
| IRSOM | 0.65 | 0.71 | 0.75 | 0.83 | 0.65 | 0.70 | 0.66 | 0.74 |
| SPECTre | 0.64 | 0.68 | - | - | 0.63 | 0.75 | 0.7 | 0.73 |

We classified the annotated ORFs of each of the four organisms as “operon” or “non-operon” (see Methods, Selection of subsets). We then calculated the AUC of precision-recall curves calculated at a overlap threshold of 70% for all five tools with either the *operon* or *non-operon* sets separately for each organism (Table 5). DeepRibo and Reparation_blast had similar or better performance for ORFs within operons (with the exception of the *Listeria* dataset). The other tools performed worse in all benchmark sets for genes located in operons compared to single-standing genes, which indicates a clear advantage of tools designed for bacteria in this regard.

Above, we found that the bacterial tools were able to detect most ORFs in example ribosomal protein and *cydAB* /*cioAB* terminal oxidase operons (Supplemental Figures S2 & S4), while IRSOM, Ribo-TISH, and SPECtre performed less well. Interestingly *cydAB* from *L. monocytogenes* overlap by 14 nt, and were detected poorly by both IRSOM and Ribo-TISH (Supplemental Figure S4D). We selected an additional, more lowly-expressed eight-gene operon (*ydjX, ydjY, ydjZ, ynjA, ynjB, ynjC, ynjD, ynjE*) in our *E. coli* dataset for inspection (Supplemental Figure S6A). Here, all genes were detected by IRSOM, and only one was missed by Reparation_blast. None of these genes were manually labeled as translated because of their overall low signal in both Ribo-seq and RNA-seq libraries. In addition, DeepRibo did not detect any of these ORFs, possibly because it has a more stringent expression cutoff. We also inspected the well-characterized overlapping ORFs *btuB* and *murI*, which share 56 bp at the 3’ end of *btuB*, in our *E. coli* dataset. Interestingly, all of the tools except Ribo-TISH called both ORFs as translated (Supplemental Figure S6B). Finally, we inspected an example of a leaderless ORF, *rluC*, in the *E. coli* dataset (Supplemental Figure S6C). Again, the same three out of the four tools detected *rluC* translation. Together, our global and single-locus observations suggest that the bacterial tools perform relatively well for both single-standing and operon-encoded genes.

#### High sensitivity comes with high false positive rate in predicting small ORFs

Genome annotations are notorious for lacking sORFs - those encoding proteins of 50 aa or less [48]. We therefore tested the performance of the tools solely on short genes by constructing a *small ORF* subset for each of the four organisms including only annotated ORFs of only 50 codons or less. The general incompleteness of sORF annotation in bacteria is supported by the *L. monocytogenes* (2.9 Mbp) and *P. aeruginosa* (6.3 Mbp) *small ORF* sets, which were smaller (7 and 12 sORFs, respectively (Table 2)) than might be expected based on their genome size compared to *E. coli* (4.6 Mbp, 114 sORFs) and *S.* Typhimurium (5.1 Mbp, 100 sORFs), which are considered some of the best annotated organisms for sORFs [21]. We therefore exclusively investigated the *E. coli* and *S.* Typhimurium *small ORF* sets, which were large enough for unbiased investigation.

Our manual labeling of the *E. coli* and *S*. Typhimurium *small ORF* subsets suggested that 55 of 114 and 31 of 100 sORFs, respectively, were translated under the investigated condition (Figure 2, top graphs and Table 2). Inspection of the tool predictions showed that DeepRibo detected 44, SPECtre 19, and Reparation_blast 19 of the 55 positively-labeled sORFs in the *E. coli small ORF* set (Supplemental Table 13). For *S.* Typhimurium, DeepRibo flagged 26 of 31 positively-labeled sORFs as translated, while Reparation_blast and SPECtre detected only nine. In contrast, IRSOM and Ribo-TISH detected hardly any of the positively-labeled sORFs in these organisms (4/3 out of 55 for *E. coli* and 5/3 out of 31 for *S.* Typhimurium), respectively. This shows that these tools do not perform well for sORF discovery in bacteria. All 19 sORFs detected by Reparation_blast in *E. coli* were also detected by DeepRibo (Figure 2). SPECtre also detected 19 sORFs, but three of these were not predicted by the other tools. Likewise, for *S.* Typhimurium, all nine sORFs called as translated by Reparation_blast were in the set detected by DeepRibo. SPECtre, which also predicted in total nine sORFs, shared one prediction with both tools and eight exclusively with DeepRibo. DeepRibo and Reparation_blast made only a few false positive predictions (seven/six of the *E. coli* sORFs not labeled as translated, and five/two for *S.* Typhimurium, respectively) and correctly predicted most of the sORFs that were labeled as not translated (52 out of 53) (Supplementary Table 13). However, SPECtre made more FP than FN predictions for both organisms (24 FP/19 TP for *E. coli* and 21 FP/9 TP for *S.* Typhimurium). Our data suggest that DeepRibo is the method of choice for sORFs, since it detects all found by Reparation_blast and almost all detected by SPECtre. Eight and five sORFs for *E. coli* and *S.* Typhimurium, respectively, were labeled as positive by manual curation but not detected by any of the tools (Figure 2).

**Figure 2:**
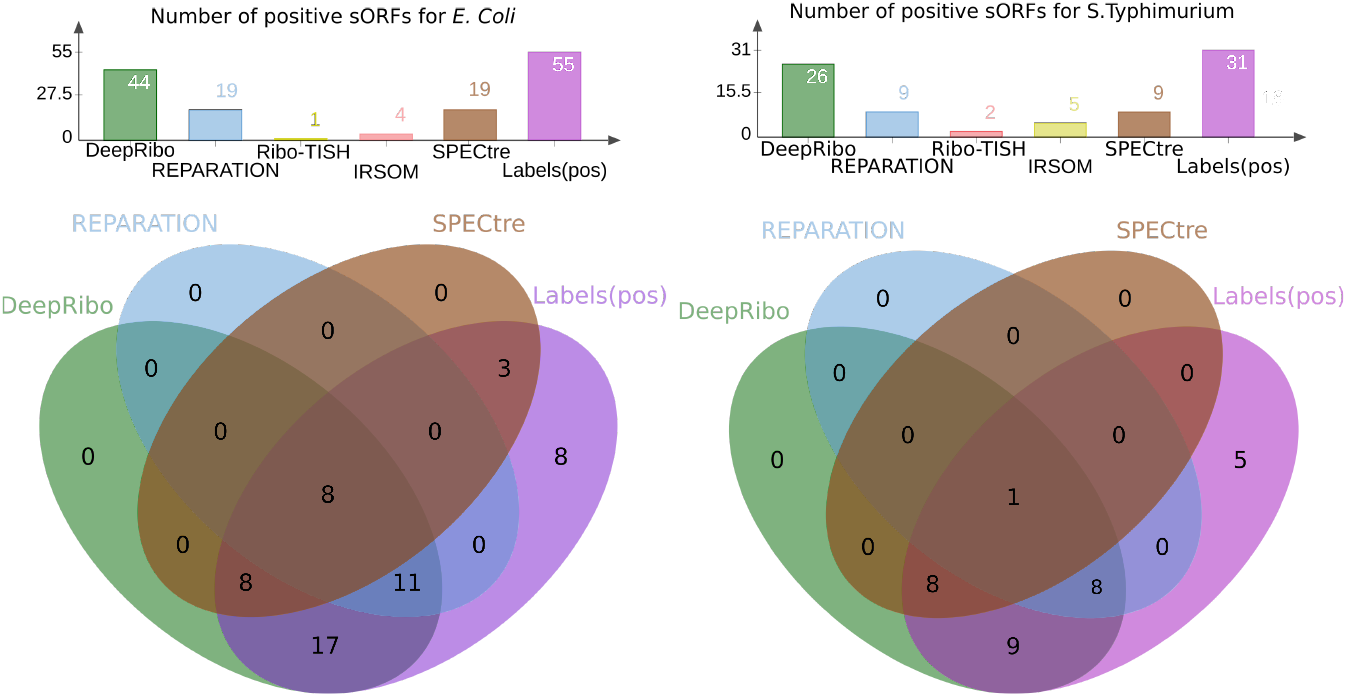
Comparison of the correctly-detected sORFs for *E. coli* and *Salmonella* Typhimurium by each tool to manual labeling. Top: the number of correctly-predicted, translated sORFs by DeepRibo, Reparation_blast, Ribo-TISH, IRSOM, SPECtre, or manual labeling. Bottom: the overlap of sORFs detected by DeepRibo (green), Reparation_blast (blue), or SPECtre (brown) with the sORFs labeled as translated (purple) for *E. coli* (left) and *S.* Typhimurium (right). The number of TP sORFs detected by the tools were determined at an overlap threshold of 70%. Ribo-TISH and IRSOM were omitted from Venn diagrams because of their low numbers of predictions.

We next inspected specific examples of positively labeled sORFs for their coverage compared to their tool predictions. The *E. coli* small membrane protein AcrZ (49 aa), a regulatory component of the AcrB-TolC antibiotic efflux pump [21] was detected by DeepRibo, Reparation_blast, and even IRSOM via RNA-seq coverage, but not Ribo-TISH (Supplemental Figure S6A). SgrT, encoded by the dual function sRNA SgrS [21], was identified as translated by DeepRibo and Reparation_blast (Supplementary Figure S6B). Again, we revisited the *cydAB* /*cioAB* operons (Supplementary Figure S4). In many proteobacteria, a small protein component of the terminal oxidase complex is encoded downstream of *cydAB* /*cioAB* [1]. For example, CydX (37 aa) of *E. coli* and *S.* Typhimurium is encoded downstream of *cydB*, while the putative sORF *cioZ* is encoded downstream of *P. aeruginosa* CioB (Supplementary Figure S4). So far, a similar small protein has not been detected in Firmicutes such as *L. monocytogenes* [1]. All three of these sORFs were manually labeled as translated in *E. coli*, *S*. Typhimurium, and *P. aeruginosa*. At an overlap threshold of 0.7, DeepRibo also detected translation of all three, while Reparation_blast only detected the enterobacterial sORFs and SPECtre detected only *E. coli cydX*. IRSOM and Ribo-TISH did not call any of the sORFs as translated. We also inspected one validated sORF from *L. monocytogenes*, since it does not encode a *cydX*. The sORF *lmo1980* [24] was labeled manually as translated and also detected only by the bacterial ORF prediction tools DeepRibo and Reparation_blast (Supplementary Figure S6C).

#### Novel *E. coli* small ORFs

Up to this point, we focused only on previously-annotated labeled ORFs. However, the discovery of novel ORFs and sORFs is one of the most interesting applications of Ribo-seq. To understand how well the different tools detect novel, potentially more challenging, sORFs, we also ran our benchmark pipeline on the plain (i.e. untreated) Ribo-seq library that was generated as part of a TIS profiling experiment to experimentally identify novel *E. coli* sORFs [54]. Of the sORFs detected by this study, 33 were verified by epitope tagging and western blotting under the same conditions as TIS profiling. We labeled the 33 sORFs based on Ribo-seq coverage alone (no RNA-seq library was available and TIS coverage was not used) without knowledge of western blot results, which suggested that 21 of the 33 sORFs showed significant Ribo-seq coverage and are likely translated. We then compared the output of DeepRibo, Reparation_blast, and Ribo-TISH to explore the overlap between the 21 positively-labeled novel sORFs and the tool predictions. We did not include SPECtre or IRSOM in this analysis, since these tools require an RNA-seq library, which was not available. However, since SPECtre did not predict any ORFs outside of the existing annotation for the other benchmark datasets (see Supplemental Tables S1, S3, and S4), this suggests it likely has a no utility in identification of novel ORFs in bacteria. Inspection of the predictions for the remaining three tools first showed that while Reparation_blast detected two of the verified sORFs, Ribo-TISH did not detect any (Supplemental Table S14). These tools were then omitted from the comparison. In total, DeepRibo predicted 20,693 potential novel ORFs of all lengths. Considering that only 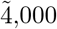 ORFs are currently annotated in *E. coli*, many of these predictions are likely false positives.

DeepRibo provides a score for each detected ORF (novel and annotated), where ORFs with a higher score are of higher confidence, to allow their ranking. However, it is left to the user to find an appropriate cutoff. Taking the top 2,000 predictions of all novel open reading frames, ranked by score, we found that DeepRibo predicted 18 of the 33 verified novel sORFs (Figure 3). To simulate the selection of novel small (≤50aa) ORFs for experimental verification we filtered for the top 100 small ORFs predicted by DeepRibo. Seven of these predicted novel sORFs (excluding *ynfU* (56aa), *yibX* (80aa)) were identified by western blots [54] (see Table 6). The next seven sORFs are then among the top 520 predictions, already a large number for evaluation by western-blot. Therefore testing of the top 100 small novel ORFs seems to be a good strategy to identify multiple novel ORFs with DeepRibo, despite missing a clear cutoff from the tool itself. However, many putative sORFs were predicted by DeepRibo with better scores than the 18/33 validated sORFs (data not shown), and many of these were not identified by TIS profiling in the original study [54]. DeepRibo predicted four novel sORFs with higher ranks than all western-blot verified sORFs, found by the original study [54]. An explanation for the different results could be that the sORFs reported by the original study were based on enrichment in the TIS library compared to the Ribo-seq library near the start codon, rather than Ribo-seq ORF coverage. This suggests that including computational sORF predictions with DeepRibo can complement TIS profiling experiments. Moreover, inclusion of approaches to utilize TIS data, which is currently not available for bacterial tools, might increase the confidence of sORF predictions by detecting those with both ORF coverage and a start codon signal.

**Table 6:** Detection of novel *E. coli* small ORFs by DeepRibo. Successfully predicted, experimentally verified novel ORFs and sORFs [54] with their score and rank for either all predictions (annotated and novel), or novel predictions only.

| Gene name | Rank all | Rank novel (all lengths) | Rank novel sORF | DeepRibo Score |
| --- | --- | --- | --- | --- |
| *yibX* | 1,784 | 7 | - | 1.372 |
| *ynfU* | 2,019 | 12 | - | 1.114 |
| *yqgG* | 2,512 | 46 | 15 | 0.169 |
| *ythB* | 2,560 | 64 | 23 | 0.006 |
| *yodE* | 2,581 | 77 | 26 | -0.006 |
| *ychT* | 2,667 | 135 | 42 | -0.464 |
| *ykiE* | 2,674 | 141 | 45 | -0.481 |
| *ymiD* | 2,680 | 145 | 46 | -0.491 |
| *yfiS* | 2,718 | 180 | 61 | -0.600 |
| *ysgD* | 2,872 | 318 | 115 | -1.115 |
| *yncP* | 3,036 | 477 | 174 | -1.464 |
| *ynaN* | 3,518 | 953 | 427 | -2.000 |
| *yadX* | 3,626 | 1,061 | 498 | -2.080 |
| *ysaE* | 3,663 | 1,098 | 519 | -2.111 |
| *yqiM* | 3,664 | 1,099 | 520 | -2.112 |
| *ybiE* | 3,930 | 1,364 | 688 | -2.282 |
| *yljB* | 4,032 | 1,466 | 759 | -2.353 |
| *ytiB* | 4,565 | 1,998 | 1,129 | -2.613 |
| *yqhJ* | 9,607 | 7,034 | 5,352 | -4.078 |

**Figure 3:**
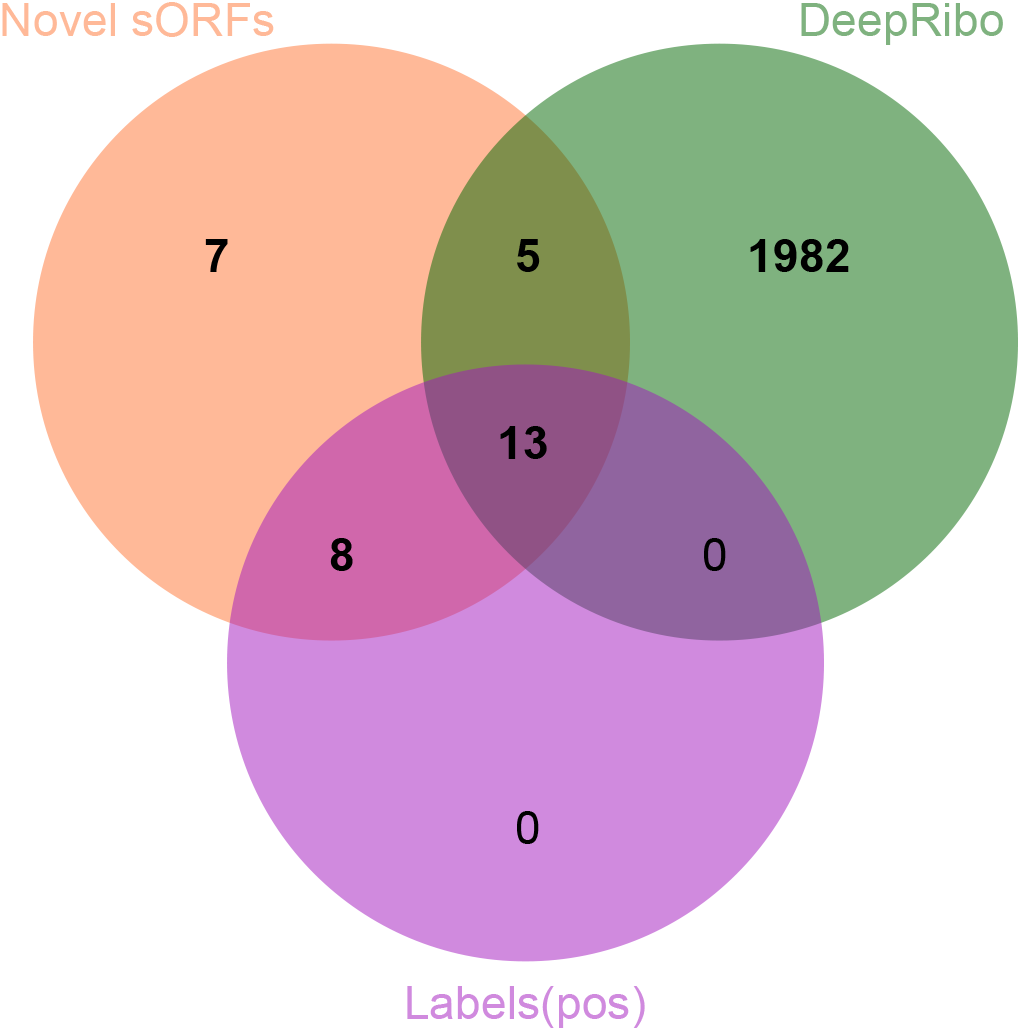
Overlap of novel sORFs detected by DeepRibo and a set of experimentally-verified sORFs in a published *E. coli* dataset. The top 2,000 predicted novel sORFs for DeepRibo were compared to 33 novel sORFs recently dtected by TIS profiling and verified by western blot in *E. coli* [54]. Also shown is the propration of the 33 sORFs labeled by manual curation of the Ribo-seq library.

**Figure 4:**
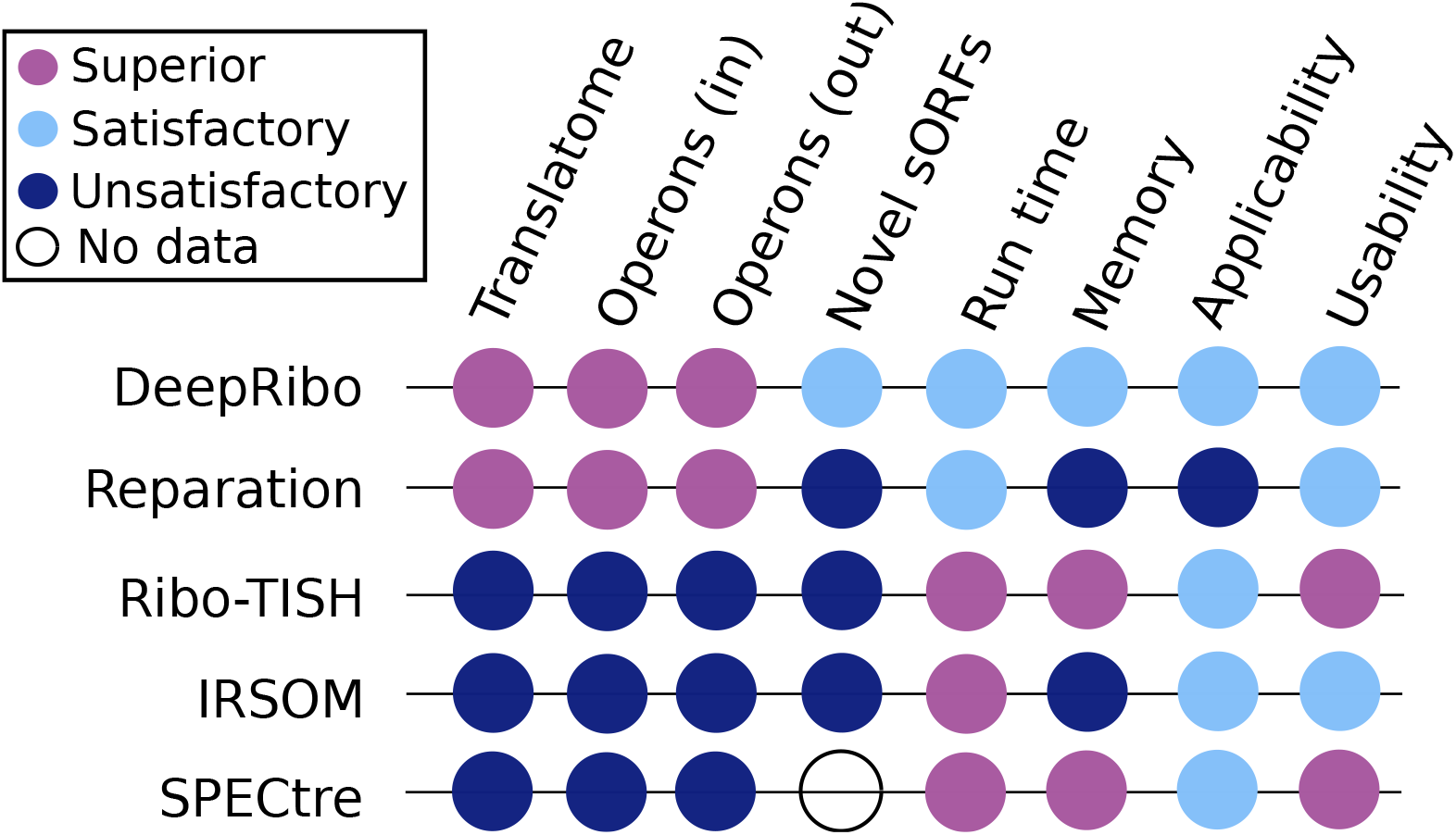
Summary of tool performance and secondary measures. Shown are the scored tool prediction performances (AUC) on the whole *translatome* and the subsets of ORFs inside and outside of operons. Furthermore, the performance of novel ORF detection outside the annotation using the results of [54] are scored. The last four columns summarize the evaluated quality measures: Here run time, memory usage, applicability for experimental design (Applicability), and user-friendliness (Usability) are scored. The evaluation system can be found in the supplement.

To provide some first insight into DeepRibo predictions vs. manual labeling, we inspected Ribo-seq coverage for several of the 33 novel sORFs. Overlap of the sORFs detected by manual labeling and DeepRibo showed 13 sORFs detected by both methods. The two sORFs with the highest DeepRibo score, *yibX* (80 aa) and *ynfU* (56 aa), showed full-read coverage in the Ribo-seq library (Supplementary Figures S7A and S7B) and both had enriched peaks in the TIS profiling library compared to the normal Ribo-seq library when 3’ end coverage was compared. Both were also manually labeled as translated. In comparison, validated *yqhJ* (19 aa) was also both labeled as translated and detected by DeepRibo. However, this candidate has the lowest DeepRibo score (−4.078) and was ranked 5352th out of all novel sORF candidates, despite having significant Ribo-seq coverage and a strongly enriched TIS peak (Supplementary Figure S7C). Other validated candidates, such as *yecV* (14 aa), were not labeled as translated in our curation or detected by DeepRibo (Supplementary Figure S7D), likely because of low Ribo-seq coverage, despite having a strongly enriched TIS peak. In contrast, *evgL* (9 aa, Supplementary Figure S8A) showed Ribo-seq coverage and a TIS signal, but was likely missed because it is below the length cutoff of DeepRibo (10 aa). Other validated sORFs not detected by DeepRibo included examples where the prediction was longer than the validated sORF (*ytgA*, 16 aa, Supplementary Figure S8B) or in a slightly different position (*yhgP*, 9 aa, Supplementary Figure S8C) possibly because the validated sORF is below the DeepRibo length cutoff.

The above observations suggest that even the bacterial prediction tools require further optimization in the context of novel sORF detection, or can be prone to missing true candidates due to expression cutoffs. However, many additional novel sORFs not reported in [54] were detected by DeepRibo with a relatively high predictive score. While some of these, such as a 44 aa sORF internal in *ftsH*, might be false positives that can be excluded based on TIS data (Supplementary Figure S8D), others could be candidates for experimental verification. Nonetheless, the ranking system of DeepRibo and the observation of the rank-distribution of verified novel candidates shows that a robust cut-off could improve the usability of DeepRibo.

### Secondary measures

Besides predictive power, other practical considerations can influence the choice of the best tool for ORF detection. We therefore also investigated quantitative (runtime and peak memory usage) and qualitative (usability, applicability) [55] secondary measures for each tool.

**Runtime and peak memory usage** of the tools were investigated in a single and multi-threading scenario. Run time and memory were analysed using the self-generated *E. coli* benchmark set. The size of the associated Ribo-seq *BAM* file is 159 MB (7,457,594 reads) and the RNA-seq *BAM* file 197 MB (9,660,815 reads). The annotation file used includes 4,379 annotated coding features. This analysis was run on a cloud instance using an AMD EPYC 7351P 16-Core processor and 128 GB of RAM, using the taskset utility for all tools.

The best runtime using only one CPU core was achieved by IRSOM, which completed analysis of the dataset in under one minute, followed by Ribo-TISH (less than 13 minutes), DeepRibo (approx. 37 minutes), and Reparation_blast (more than 2 hours) (Table 7). When running DeepRibo, we noticed that it ran reproducibly faster on a single thread compared to multiple threads. In addition, DeepRibo ignored the maximum number of threads assigned via command-line attribute if the maximum number of cores was not restricted by the operating system. This behaviour was reproduced on another cloud instance with a different hardware setup. SPECtre had an average runtime compared to the other tools. We did not observe a difference in runtime when providing multiple cores when using the default settings of SPECtre. On a single core, Ribo-TISH had the lowest peak memory consumption (119 MB), followed by SPECtre (1,494 MB), DeepRibo (3,977 MB), Reparation_blast (5,517 MB), and IRSOM (8,027 MB).

**Table 7:**
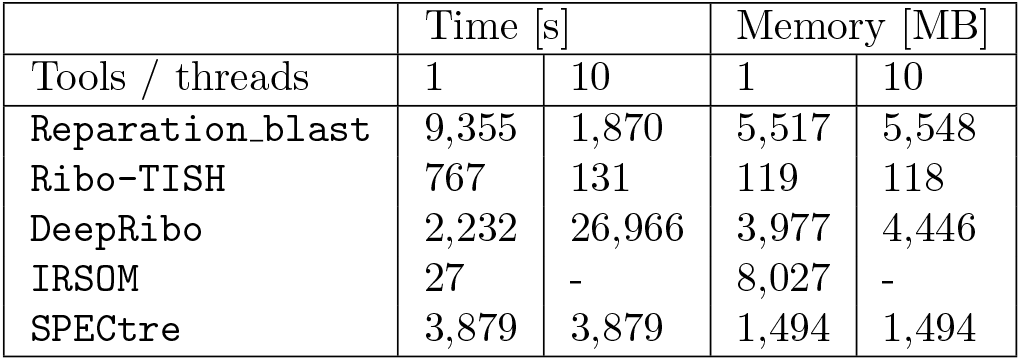
Runtime and peak memory consumption. Runtime and peak memory consumption for each tool running on a virtual machine with either one or ten CPU cores, processing one library of the self-generated *E. coli* dataset.

**Applicability** of a tool can also contribute to its suitability for a specific task. Ribo-TISH is the only tool out of the five tested that supports the input of replicates. Reparation_blast, on the other hand, was the only tool that does not produce a deterministic output, meaning that the results of the tool with identical inputs are different between calls. Only DeepRibo uses a standard output format (*BEDGRAPH*) and also only via an included post-processing script, while the output of the other tools has to be parsed or converted for downstream analysis by the user. None of the tools uses unit testing, to ascertain the correctness of functions and the reliability of results. Nevertheless, the results of all tools were consistent over different species and annotations. We scored the applicability of the tools as detailed in the supplemental material (Subsection D.6, Applicability).

**Usability** determines how user-friendly a tool is. We scored the usability of each tool as detailed in the supplemental material (Subsection D.7, Usability). The five benchmarked tools are stably available from software hosting platforms. Only Ribo-TISH and Reparation_blast can be installed with dependencies via a package management system (and only because we included them). With the exception of Ribo-TISH, all tools have a sample dataset available for testing. DeepRibo and Ribo-TISH feature change-logs. They also feature, as well as SPECtre, a versioning scheme - a key criterion for reproducibility. The documentation of the tools has varying levels of detail and completeness, but all have documented tool dependencies. However, the command line parameters of IRSOM are not documented, DeepRibo is missing documentation concerning its required input, and the output documentation of IRSOM as well as REPARATION is either missing or difficult to find. The published version of SPECtre accepts only ensembl-formatted *GTF* annotation input, which will make it necessary for many users to specifically preprocess their annotation. All tools are open source, including REPARATION in the Reparation_blast variant.

## 4 CONCLUSIONS

With RiboReport, we aimed to identify the best available tools for Ribo-seq based ORF detection in bacteria using a set of trusted ORFs that we have generated from datasets generated in diverse species. Astoundingly, out of the nine tools found in literature, only four (DeepRibo, Reparation_blast, SPECtre, and Ribo-TISH) were compatible with bacterial annotations and genomes (Table 3). In addition, the coding potential detection tool IRSOM, which uses only transcriptome data, was added to investigate the performance gain achieved by using Ribo-seq data together with specialized ORF detection tools. While the predictive performance of DeepRibo and Reparation_blast was superior to the other tools, their runtime and peak memory consumption were substantially higher than for IRSOM, SPECtre and Ribo-TISH. None of the tools was available via a package manager.

DeepRibo and Reparation_blast showed a superior predictive performance over SPECtre, Ribo-TISH, and IRSOM for all organisms and all annotated ORF sets (*translatome*, *small ORFs*, *operon*, and *non-operon*). A set of recently identified and validated sORFs outside of the *E. coli* annotation [54] was used to test novel sORF detection. These sORFs were poorly detected by all tools, with the exception of DeepRibo. It predicted 18 of the 33 sORFs, but most of these predictions did not have a high rank (Table 6).

The high sensitivity of DeepRibo appears to come at a cost of a high false positive rate. While a metric is generated by the tool to provide a way to sort for higher confidence candidates, a robust cutoff is not offered for this score to allow investigation of strong candidates only. However testing of the top 100 small novel ORFs seems to be a strategy to identify some novel ORFs.

For the tools that we could not test, there was no mentioning of their taxonomic scope or if they are applicable beyond the scope of what they had been designed and tested on. Ribo-TISH, while unsatisfactory in terms of predictive power, was also clearly not designed with bacterial data in mind. However, it was the only tool that supports replicates as input, as well as TIS Ribo-seq data. As TIS profiling is now established in bacteria and archaea [17, 34, 54], we expect this to be an essential capability of future tools. Looking to the future, we hope that support for TIS data, replicates, and non-standard organisms is considered in new tools or improved versions of the current tools.

## 5 Competing interests

There is NO Competing Interest.

## 6 Author contributions statement

F.E., R.G., S.L.S., T.M., and R.B. designed the study; S.L.S performed the experiments; F.E., S.L.S., R.G, and T.M. screened databases for bacterial Ribo-Seq data sets; S.L.S. performed manual labeling of the translated regions; F.E. retrieved and processed the proteomics data; O.A. computed the operon regions; R.G performed high throughput sequencing analysis, tool testing, and ORF predictions; T.M. performed the benchmark analysis and created the benchmark plots; R.B. and C.S. provided the funding, and all authors jointly wrote the manuscript.

## 7 Acknowledgments

We thank the members of DFG SPP 2002 (“Small Proteins in Prokaryotes: An Unexplored World”) for constructive discussions, Thorsten Bischler for assistance with data analysis, and Ann-Janine Imsiecke for assistance with Ribo-seq.

## 8 Funding

This work was supported by the German Research Foundation (DFG), Germany’s Excellence Strategy (CIBSS - EXC-2189 - Project ID 390939984) and DFG SCHM 2663/3, by the High Performance and Cloud Computing Group, University Tübingen via bwHPC, the BMBF-funded de.NBI Cloud within the German Network for Bioinformatics Infrastructure (de.NBI) (031A532B, 031A533A, 031A533B, 031A534A, 031A535A, 031A537A, 031A537B, 031A537C, 031A537D, 031A538A), DFG INST 37/935-1 FUGG.R.G., DFG grants BA 2168/21-1 SPP 2002 Small Proteins in Prokaryotes: An Unexplored World; BA 2168/23-1 SPP 2141 Z-Projekt CRISPR Bioinformatik; BMBF “RNAProNet - 031L0164B, BMBF grant 031A538A de.NBI; DFG grants to B.A SH580/7-1 and Sh580/8-1 within the DFG SPP2002 to C.M.S.”

## Notes

https://github.com/RickGelhausen/RiboReport

### Competing Interest Statement

The authors have declared no competing interest.

