## Supplemental material for "RiboReport - Benchmarking tools for ribosome profiling-based identification of open reading frames in bacteria"

### RiboReport - Supplemental Material

#### A Introduction

This document contains supplemental material for *RiboReport - Benchmarking tools for ribosome profiling-based identification of open reading frames in bacteria.*

#### B Validation of labeling method

We validated our labeling approach by comparison to available published MS datasets (proteomics) for the same strains grown under similar conditions. Comparing the translated ORFs found by our labeling method to those detected in the retrieved proteomics data for each organism showed that the majority of genes labeled as translated based on Ribo-seq were also detected by MS (Supplemental Figure S1)(on average 83.81% across all four organisms), thereby validating our labeling procedure.

To further showcase the overlap of human labeling in the genome browser versus MS to generate a robust benchmark set, we selected a set of conserved and highly translated ORFs that could be compared in all four organisms: a long ribosomal protein island that is highly conserved in bacteria (genes *rpmJ* to *secY*) and also features several sORFs with less than 50 aa. The genome browser tracks show that for *E. coli*, *P. aeruginosa*, and *L. monocytogenes*, both MS and labeling detected all 22 of these ORFs and that the entire island showed strong

Ribo-seq coverage (Figs S2A - S2C). In contrast, while all 22 ORFs were labeled as translated in *S.* Typhimurium only five were detected by MS, consistent with the generally lower sensitivity of the *S.* Typhimurium MS dataset (Figure S1).

As an additional check of our labeling procedure, we also labeled a selection of tRNAs and annotated non-coding RNAs based on our *de novo E. coli* dataset (data not shown). None of the tRNAs examined (86) were called as translated based on comparison of Ribo-seq and RNA-seq coverage. Several well-characterized ncRNAs were also labeled as not translated, such as the regulatory small RNAs MicA (Supplemental Figure S3A), Spot 42, SdsR, and GlmY, as well as the housekeeping RNA components of the signal recognition particle (SRP RNA) and RNase P (RnpB). However, some known non-coding RNAs were labeled as translated, such as CsrC (Supplemental Figure S3B) and tmRNA (data not shown). Some of these false positives could be due to resistance to RNase digestion or association with ribosomes (tmRNA). However, some, such as the annotated non-coding RNA RyeG, were recently shown to encode translated sORFs (Supplemental Figure S3C, *yodE*, 48 aa) [1]. We also inspected an ORF that was detected by MS, but not labeled as translated, *katE*. This ORF was not labeled as translated because of low coverage (generally below 10 relative reads) (Supplemental Figure S3D). Its detection by MS suggests either the conditions for Ribo-seq and MS were slightly different, or that KatE is a relatively stable protein that is translated at an earlier growth phase from an mRNA that is not highly expressed during exponential growth. Nonetheless, while these examples (Supplemental Figure S3B and S3D) highlight some limitations of our labeling approach, overall the overlap of labels and MS, together with accurate labeling of known non-coding RNAs and sORFs, demonstrates the validity of our approach.

#### C Supplementary Result Figures

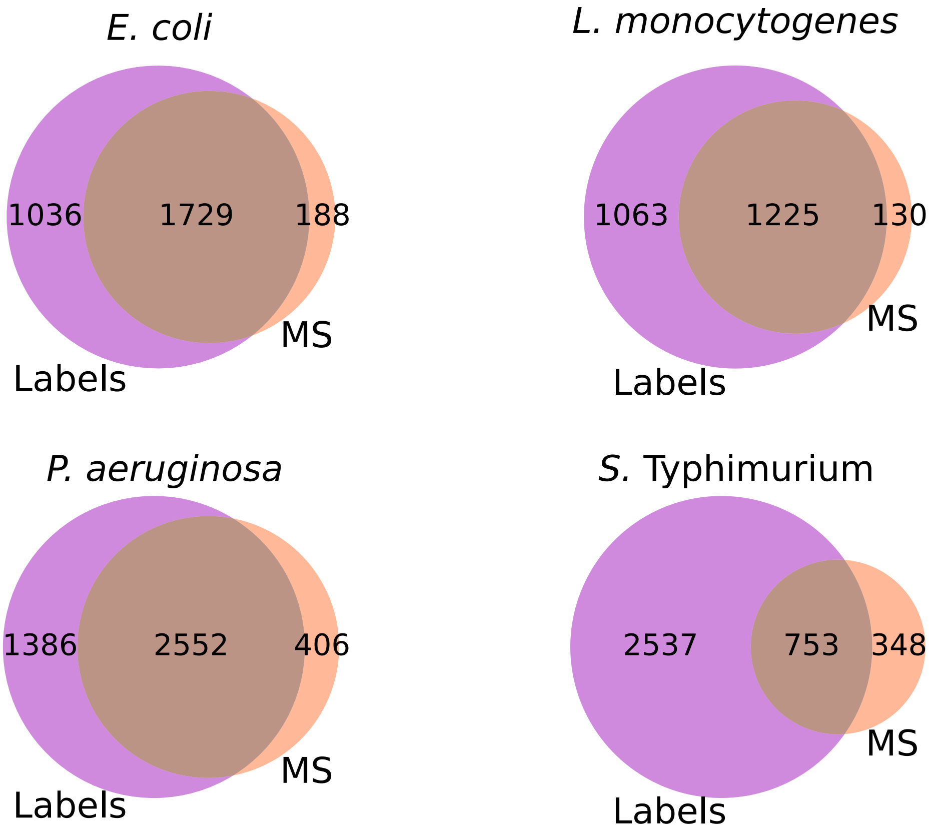

**Fig. S1: Comparison between ORFs labeled as translated with Ribo-seq data and ORFs detected with proteomics** The intersection between annotated ORFs labeled as translated based on Ribo-seq data (Labels) and those detected by mass spectrometry (MS) for the four benchmark datasets. The majority of the genes detected by mass spectrometry are also present in the labeled benchmark dataset. The percent overlapping MS ORFs which are also found by our labeling method, for each organism are as follows: *E. coli*, 90%; *L. monocytogenes*, 90%; *P. aeruginosa*, 86%; *S.* Typhimurium, 68%; and on average 83.8%. The large overlap validates the manual labeling strategy we employed to create the benchmark datasets.

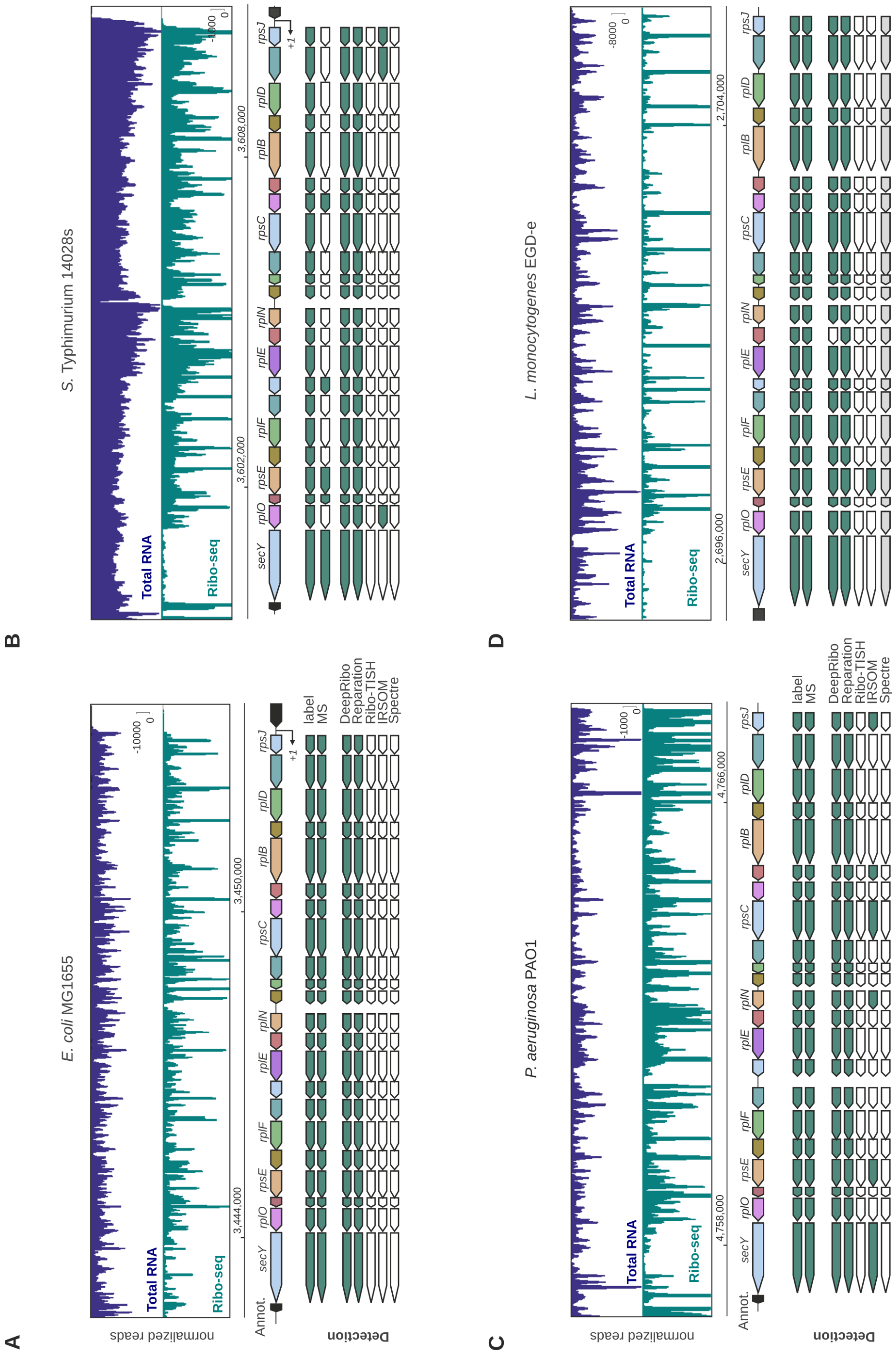

**Fig. S2:** Comparison of manual Ribo-seq curation (labels) and proteomics data (MS) for representative genes for the *translatome* sets of the four organisms. The conserved and syntenic region between *rpmJ* and *secY*, which includes several highly expressed and essential ORFs of different lengths, was compared for *E. coli* **(A)**, *S.* Typhimurium **(B)** *P. aeruginosa* **(C)**, and *L. monocytogenes* **(D)**. Corresponding genes between the organisms are labeled with the same colour in the annotation. Genes that are detected in the recovered proteomics dataset (MS), by manual curation of the Ribo-seq data (Label), or by the indicated tools with the RNA-seq or Ribo-seq libraries are indicated in green. Those that are not detected by MS or labeled as not translated are white. Arrows in grey indicate that predictions could not be generated for the genome/dataset. Transcriptional start sites, if available, are indicated with +1.

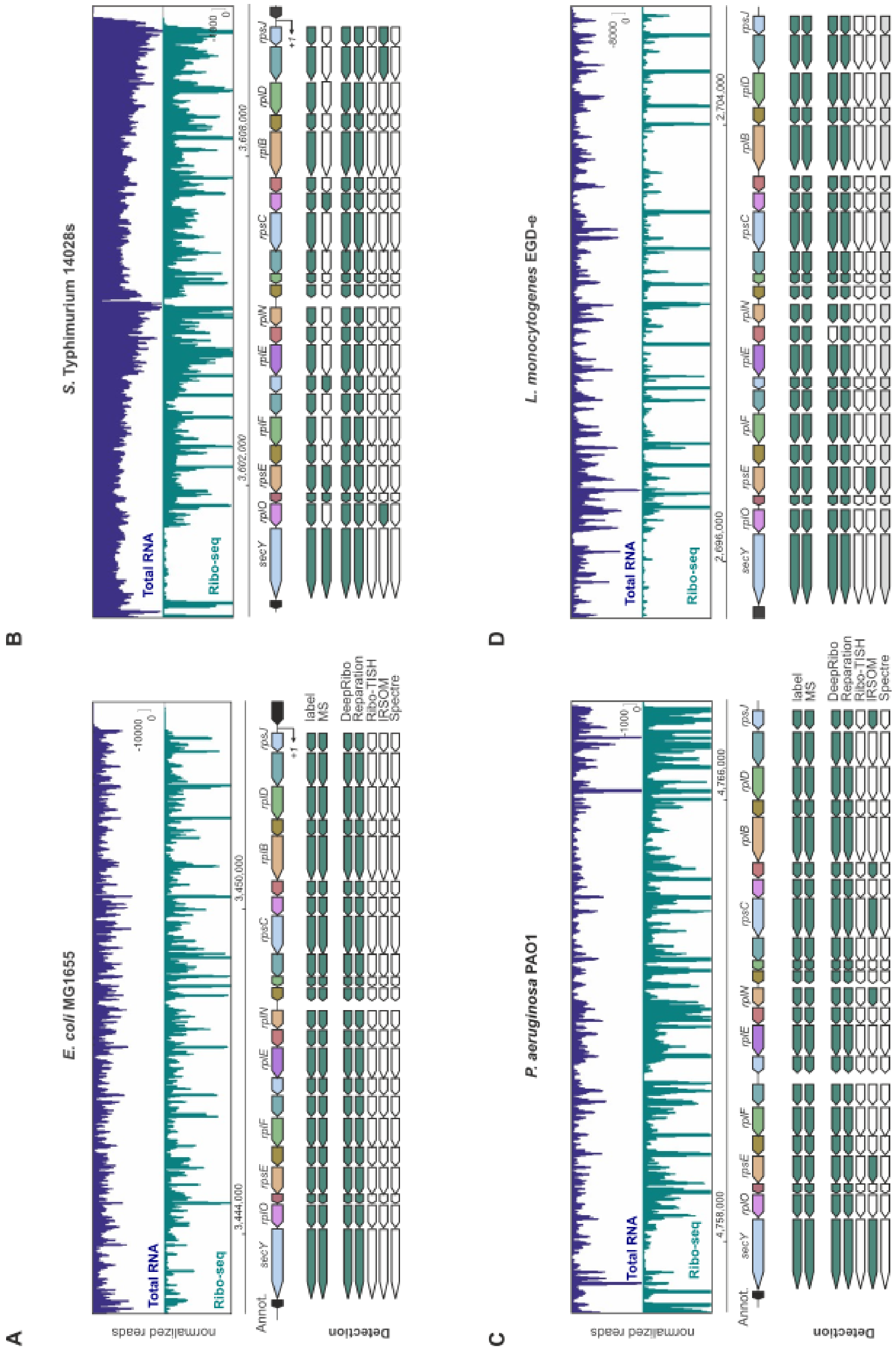
A
B
C
D
E. coli MG1655
S. Typhimurium 14028s
P. aeruginosa PAO1
L. monocytogenes EGD-e
normalized reads
Total RNA
Ribo-seq
Annot.
Detection
label
MS
DeepRibo
Reparation
Ribo-TISH
IRSOM
Spectre
secY
rplO
rpsE
rplF
rplE
rplN
rpsC
rplB
rplD
rpsJ

**Fig. S3**

**Fig. S3** ***(previous page)*****:** Assessing the quality/accuracy of manual curation for well-characterized and/or validated *E. coli* genes. An example of a true negative, false positive, true positive, and false negative manual curation from the *de novo E. coli* dataset were selected. **(A)** The non-coding base-pairing regulatory RNA MicA was correctly labeled as not translated based on Ribo-seq coverage and is a true-negative. **(B)** The non-coding RNA CsrC shows coverage in the Ribo-seq library and was false-positively labeled as translated. **(C)** The newly validated sORF true-positive *yodE* (48 aa) [2], encoded on the annotated RyeG sRNA, was labeled as translated based on Ribo-seq. **(D)** The *katE* ORF was not labeled as translated and shows low coverage in the RNA-seq and Ribo-seq libraries, but has been detected by MS under similar growth conditions. Tool predictions and MS data for annotated coding genes are also included for comparison. Detection (green arrows) or no detection (white arrows) was determined at a 70% overlap threshold.

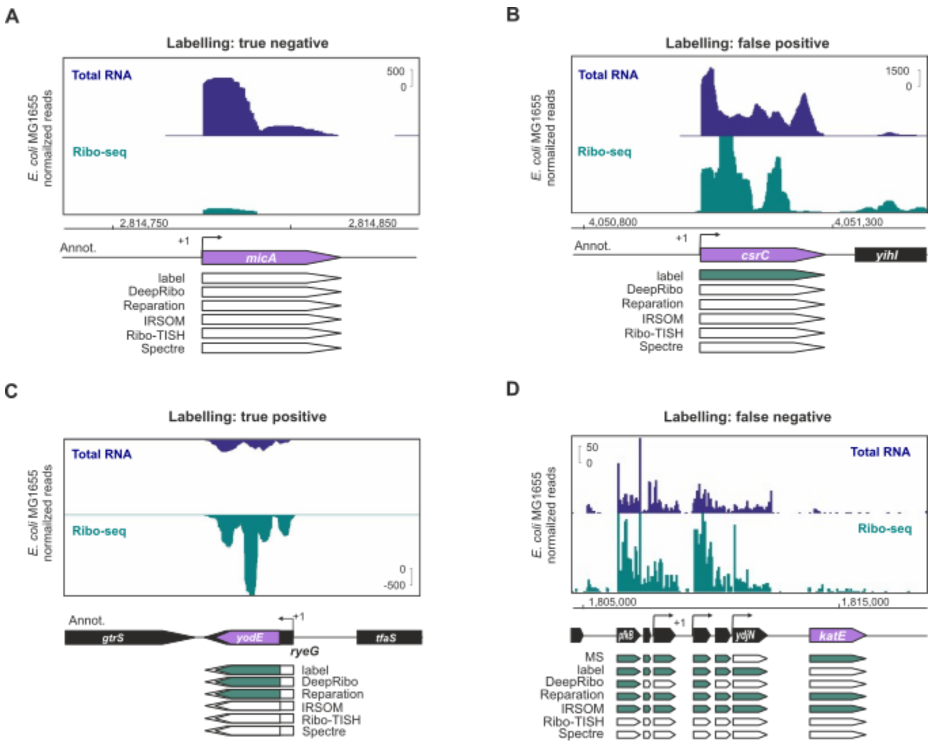

**Fig. S4:** Comparison of detection of corresponding ORFs from the *translatome* and *small ORFs* sets by the four tested tools for **(A)** *E. coli*, **(B)** *S.* Typhimurium, **(C)** *P. aeruginosa*, and **(D)** *L. monocytogenes*. The ORFs annotated in syntenic *cydAB*/*cioAB* regions, encoding terminal oxidase complexes, were inspected for MS detection, detection based on Ribo-seq by either manual labeling or the computational tools `DeepRibo`, `REPARATION_blast`, `SPECtre`, or `Ribo-TISH`), or from RNA-seq coverage (`IRSOM`). Corresponding genes between the organisms are labeled in the same colour in the annotation track. Genes encoding associated small protein components of the complexes (pink arrows) are annotated downstream of *cydAB* so far only in *E. coli*/*S.* Typhimurium (*cydX*) and *P. aeruginosa* (*cydZ*) but have so far not been detected in the *L. monocytogenes cydAB* operon [3]. Detected/labeled ORFs are green, while those not detected are white. Genes in grey indicate that predictions could not be generated for the genome/dataset. Detection (green arrows) or no detection (white arrows) was determined at a 70% overlap threshold. Spaces between *cydA* and *cydB* are as follows: *E. coli*: 15 nt, *S.* Typhimurium: 15 nt, *P. aeruginosa*: 3 nt, and *L. monocytogenes* (*cioA*,*cioB*), 14 nt overlap. For all panels, detection (green arrows) or no detection (white arrows) was determined at a 70% overlap threshold. Arrows in grey indicate that predictions could not be generated for the genome/dataset (`SPECtre`).

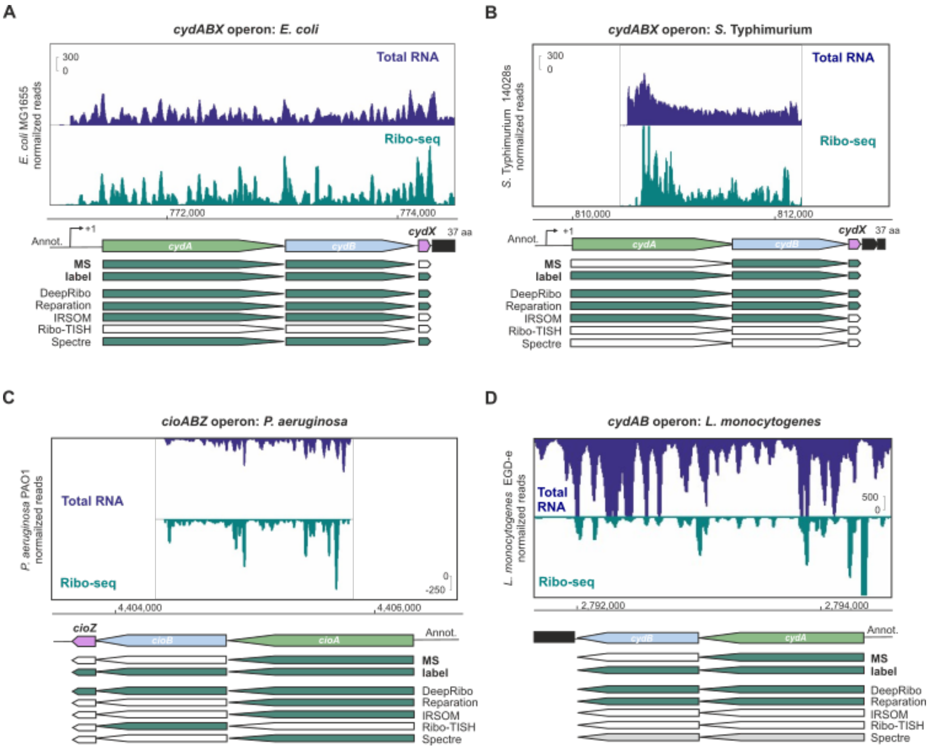

**Fig. S5:** Detection of examples of ORFs in bacteria-specific contexts in *E. coli*. **(A)** A set of eight lowly-transcribed polycistronic ORFs (*ydjX*-*ynjE*) was not labeled as translated based on Ribo-seq because of low overall coverage, but protein products from three of these ORFs were detected by MS under similar growth conditions. In contrast, `SPECtre` detected three, `REPARATION_blast` seven, and `IRSOM` eight ORFs, respectively. None of the ORFs were detected by `DeepRibo`. **(B)** Comparison of the performance of the tools on overlapping genes. The overlapping *btuB*-*murI* operon was selected from the literature and inspected. Both ORFs were detected in the MS dataset, labeled as translated based on Ribo-seq, and were also identified by `DeepRibo`, `REPARATION_blast`, and `IRSOM`. **(C)** The leaderless ORF *rluC*, which was detected by MS and labeled based on Ribo-seq, was also detected by the prediction tools `DeepRibo`, `REPARATION_blast`, and `IRSOM`, even without a Shine-Dalgarno sequence. For all panels, detection (green arrows) or no detection (white arrows) was determined at a 70% overlap threshold.

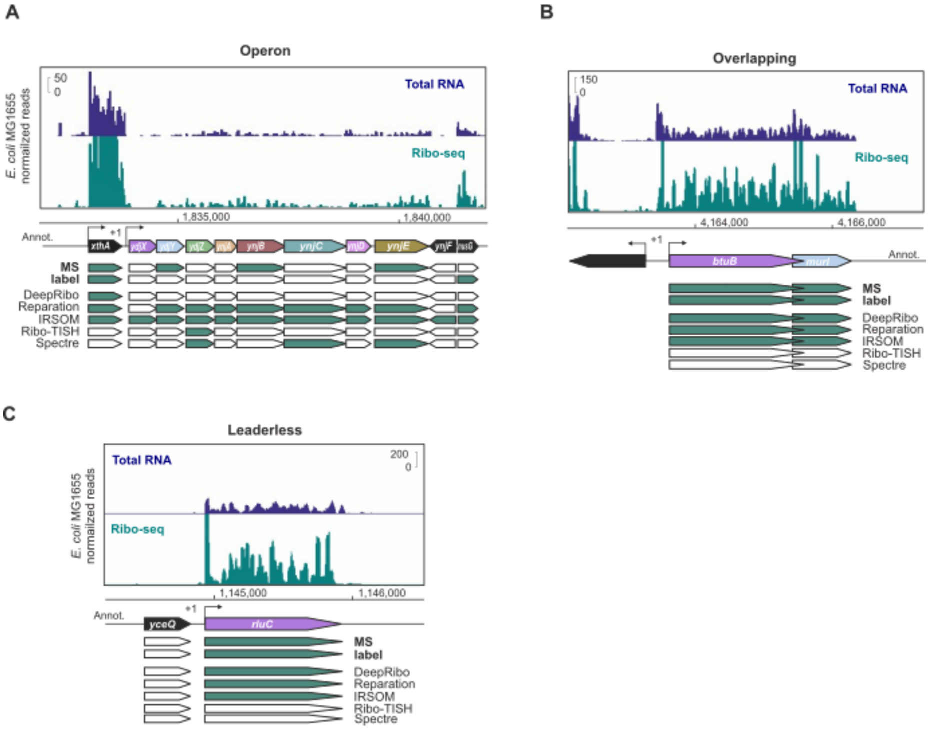

**Fig. S6:** Detection of annotated sORFs. **(A)** The sORF *acrZ* (49 aa, [4]) was labeled as translated by manual curation of Ribo-seq coverage but was not detected in the MS dataset. Translation of *acrZ* was also called as translated by `DeepRibo`, `REPARATION_blast`, and `IRSOM`. **(B)** The dual function RNA SgrS acts as both a base-pairing repressor, and also encodes the small protein SgrT (43 aa, [5]). SgrT was not detected by MS under these conditions but was labeled as translated based on Ribo-seq coverage and was detected by `DeepRibo` and `REPARATION_blast`. **(C)** The validated *L. monocytogenes* sORF *lmo1980* (45 aa) was not detected by MS but was detected by the bacterial tools `DeepRibo` and `REPARATION_blast`. For all panels, detection (green arrows) or no detection (white arrows) was determined at a 70% overlap threshold. Arrows in grey **(C)** indicate that predictions could not be generated for the genome/dataset.

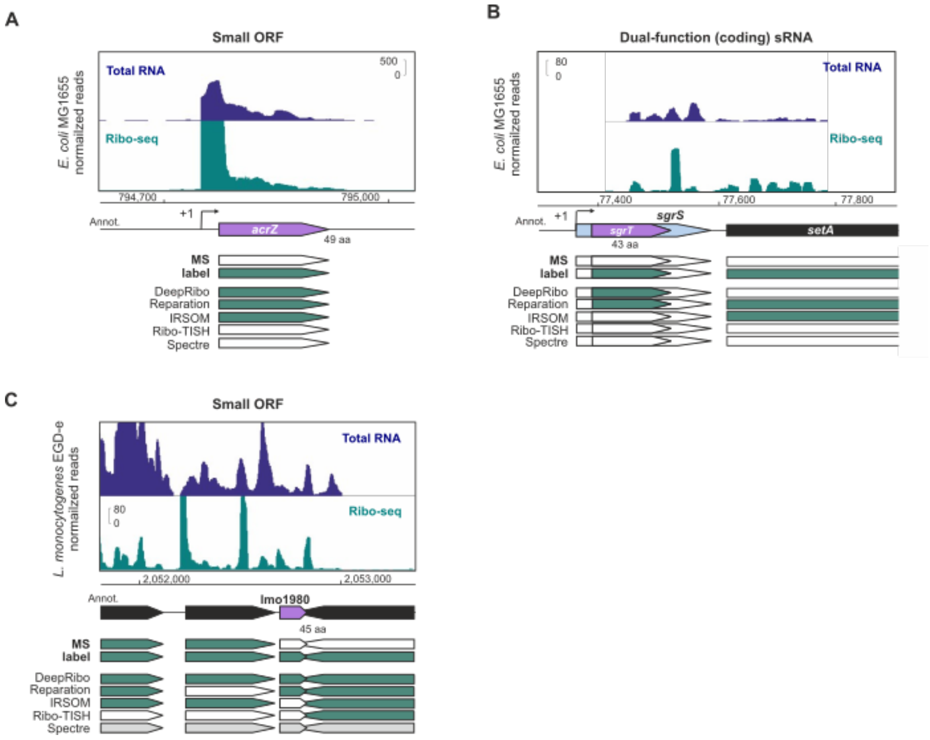

**Fig. S7:** Detection of novel, western blot-validated *E. coli* sORFs first discovered by translation initiation site (TIS) profiling based on Ribo-seq coverage by `DeepRibo` or manual labeling. The translation of 33 western blot validated sORFs [2] was labeled by manual curation of the same Ribo-seq data, and compared to `DeepRibo` predictions at an overlap threshold of 70%. It utilized the Ribo-seq data shown on top, while the study also utilized the TIS-track. **(A)** The antisense sORF *yibX* (80 aa) was the top-ranked validated sORF predicted by `DeepRibo`. The partially overlapping validated sORF *yibX-S* (24 aa) was not detected by `DeepRibo`. **(B)** The second-highest ranked sORF *ynfU* (56 aa). **(C)** The lowest-ranked detected sORF *yqhJ* (19 aa). **(D)** The validated sORF *yecV* (14 aa) was not manually labeled as translated or predicted by `DeepRibo` based on normal Ribo-seq coverage, but shows a strongly enriched TIS peak. For all panels, ORFs in green were detected as translated by [2], manual labeling, or `DeepRibo` as indicated on the left. White ORFs were not detected. The TIS track was not used for manual curation or `DeepRibo` predictions and was only included for screenshots.

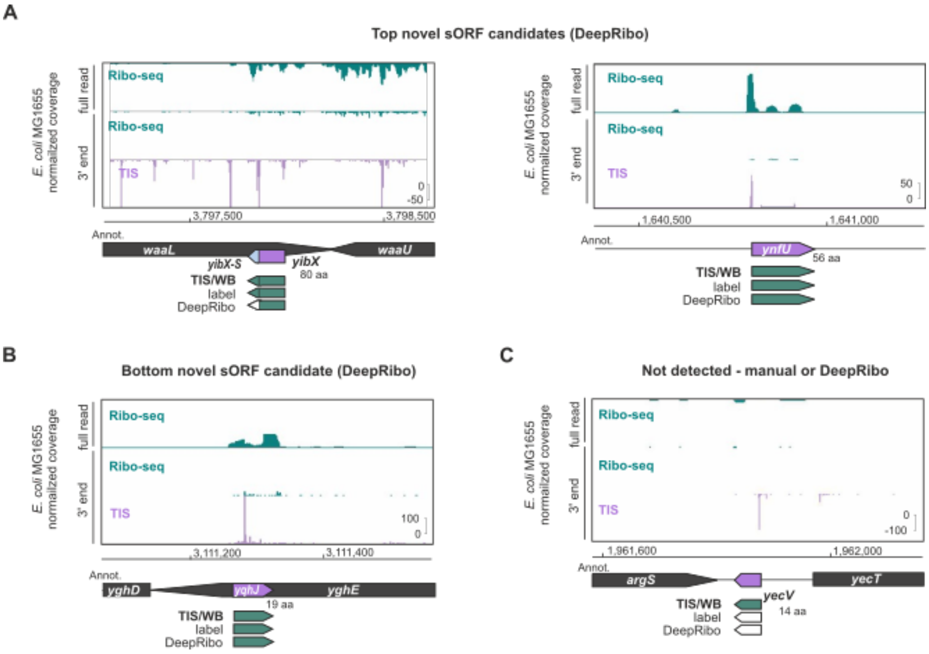

**Fig. S8:** Detection of novel, western blot-validated *E. coli* sORFs first discovered by translation initiation site (TIS) profiling based on Ribo-seq coverage by `DeepRibo` or manual labeling. The translation of 33 western blot-validated sORFs previously identified and validated in *E. coli*[2] was labeled by manual curation of the same Ribo-seq data. Labels were compared to `DeepRibo` predictions at an overlap threshold of 70%. **(A)** A validated sORF overlapping the $5'$ UTR and $5'$ end of *evgA* (*evgL*, 9 aa) was labeled as translated, but not detected by `DeepRibo`, possibly because it is below the length cutoff of the tool. **(B)** The sORF *ytgA* (16 aa) in the 5'UTR of *lptF* was labeled as translated, but the `DeepRibo` prediction is extended at the $5'$ end by 3 codons. **(C)** A 13 aa sORF was called by `DeepRibo` in the region of the validated sORF *yhgP* (9 aa), which is shorter than the `DeepRibo` length cutoff. **(D)** A novel sORF (NC_000913.3:3325012-3325146:-) that was not reported in Weaver *et al.*, but was the top-ranked sORF predicted by `DeepRibo`, at the $3'$ end of *ftsH*. No TIS peak is visible for this sORF.For all panels, ORFs in green were detected as translated by [2], manual labeling, or `DeepRibo` as indicated on the left. White ORFs were not detected. The TIS track was not used for manual curation or `DeepRibo` predictions and was only included for screenshots.

### D Supplementary Result Tables

#### D.1 Translatome

**Table S1:** Statistical evaluation of tool performance for *translatome* ORFs for the *E. coli* benchmark dataset using overlap thresholds of 0.01, 0.7, or 0.9. The overlap threshold is the percent of the ORF length that must overlap with the prediction. The overlap threshold needs to be fulfilled both for prediction with labeled ORF and vice versa. The true positives (TP), true negatives (TN), false positives (FP), and false negatives (FN) are based on the numbers of ORFs or predictions given the current overlap threshold. The sensitivity or TP rate (TPR), specificity or TN rate (TNR), FN rate (FNR), precision or positive predictive value (PPV), false discovery rate (FDR), F1 measure, and the accuracy are calculated based on the given TP, TN, FP, and FN values. Since more than one prediction meeting the specified overlap threshold can overlap with one ORF, the number of additional overlapping predictions are reported in the suboptimal TP (sTP) and suboptimal FP (sFP) columns. The number of prediction which do not overlap, given the overlap cutoff, with any annotated gene are displayed int the column no gene found (no_gene)

| TP | TN | FP | FN | TPR | TNR | FNR | PPV | FDR | F1 | accuracy | sTP | sFP | no_gene | tool |
|---|---|---|---|---|---|---|---|---|---|---|---|---|---|---|
| ***E. coli*** **and using overlap of 0.01** | | | | | | | | | | | | | | |
| 2708 | 640 | 843 | 57 | 0.98 | 0.43 | 0.02 | 0.76 | 0.24 | 0.86 | 0.79 | 55 | 10 | 145 | `REPARATION_blast` |
| 1750 | 1214 | 269 | 1015 | 0.63 | 0.82 | 0.37 | 0.87 | 0.13 | 0.73 | 0.70 | 2571 | 157 | 72 | `Ribo-TISH` |
| 2670 | 1203 | 280 | 95 | 0.97 | 0.81 | 0.03 | 0.91 | 0.09 | 0.93 | 0.91 | 16315 | 342 | 2536 | `DeepRibo` |
| 1452 | 755 | 728 | 1313 | 0.53 | 0.51 | 0.47 | 0.67 | 0.33 | 0.59 | 0.52 | 51 | 33 | 185 | `IRSOM` |
| 1088 | 809 | 674 | 1677 | 0.39 | 0.55 | 0.61 | 0.62 | 0.38 | 0.48 | 0.45 | 0 | 0 | 0 | `SPECtre` |
| ***E. coli*** **and using overlap of 0.7** | | | | | | | | | | | | | | |
| 2697 | 712 | 771 | 68 | 0.98 | 0.48 | 0.02 | 0.78 | 0.22 | 0.87 | 0.80 | 1 | 0 | 275 | `REPARATION_blast` |
| 84 | 1414 | 69 | 2681 | 0.03 | 0.95 | 0.97 | 0.55 | 0.45 | 0.06 | 0.35 | 59 | 26 | 4579 | `Ribo-TISH` |
| 2305 | 1435 | 48 | 460 | 0.83 | 0.97 | 0.17 | 0.98 | 0.02 | 0.90 | 0.88 | 17 | 1 | 19748 | `DeepRibo` |
| 1408 | 782 | 701 | 1357 | 0.51 | 0.53 | 0.49 | 0.67 | 0.33 | 0.58 | 0.52 | 6 | 7 | 309 | `IRSOM` |
| 1087 | 812 | 671 | 1678 | 0.39 | 0.55 | 0.61 | 0.62 | 0.38 | 0.48 | 0.45 | 0 | 0 | 0 | `SPECtre` |
| ***E. coli*** **and using overlap of 0.9** | | | | | | | | | | | | | | |
| 2656 | 766 | 717 | 109 | 0.96 | 0.52 | 0.04 | 0.79 | 0.21 | 0.87 | 0.81 | 0 | 0 | 371 | `REPARATION_blast` |
| 51 | 1442 | 41 | 2714 | 0.02 | 0.97 | 0.98 | 0.55 | 0.45 | 0.04 | 0.35 | 9 | 4 | 4712 | `Ribo-TISH` |
| 2246 | 1438 | 45 | 519 | 0.81 | 0.97 | 0.19 | 0.98 | 0.02 | 0.89 | 0.87 | 3 | 0 | 19825 | `DeepRibo` |
| 1408 | 782 | 701 | 1357 | 0.51 | 0.53 | 0.49 | 0.67 | 0.33 | 0.58 | 0.52 | 2 | 2 | 318 | `IRSOM` |
| 1087 | 812 | 671 | 1678 | 0.39 | 0.55 | 0.61 | 0.62 | 0.38 | 0.48 | 0.45 | 0 | 0 | 0 | `SPECtre` |

**Table S2:** Statistical evaluation of the *translatome* ORFs for the benchmark dataset *L. monocytogenes* using overlap thresholds of 0.01, 0.7, or 0.9. A detailed column description can be found in main Table S1.

| TP | TN | FP | FN | TPR | TNR | FNR | PPV | FDR | F1 | accuracy | sTP | sFP | no_gene | tool |
|---|---|---|---|---|---|---|---|---|---|---|---|---|---|---|
| ***L. monocytogenes*** **and using overlap of 0.01** | | | | | | | | | | | | | | |
| 1913 | 349 | 230 | 375 | 0.84 | 0.60 | 0.16 | 0.89 | 0.11 | 0.86 | 0.79 | 48 | 4 | 44 | `REPARATION_blast` |
| 1023 | 390 | 189 | 1265 | 0.45 | 0.67 | 0.55 | 0.84 | 0.16 | 0.58 | 0.49 | 695 | 110 | 82 | `Ribo-TISH` |
| 2284 | 56 | 523 | 4 | 1.00 | 0.10 | 0.00 | 0.81 | 0.19 | 0.90 | 0.82 | 15433 | 2051 | 4279 | `DeepRibo` |
| 1099 | 248 | 331 | 1189 | 0.48 | 0.43 | 0.52 | 0.77 | 0.23 | 0.59 | 0.47 | 117 | 52 | 107 | `IRSOM` |
| - | - | - | - | - | - | - | - | - | - | - | - | - | - | `SPECtre` |
| ***L. monocytogenes*** **and using overlap of 0.7** | | | | | | | | | | | | | | |
| 1843 | 378 | 201 | 445 | 0.81 | 0.65 | 0.19 | 0.90 | 0.10 | 0.85 | 0.77 | 0 | 0 | 176 | `REPARATION_blast` |
| 55 | 556 | 23 | 2233 | 0.02 | 0.96 | 0.98 | 0.71 | 0.29 | 0.05 | 0.21 | 32 | 18 | 1969 | `Ribo-TISH` |
| 2194 | 215 | 364 | 94 | 0.96 | 0.37 | 0.04 | 0.86 | 0.14 | 0.91 | 0.84 | 4 | 1 | 21932 | `DeepRibo` |
| 975 | 289 | 290 | 1313 | 0.43 | 0.50 | 0.57 | 0.77 | 0.23 | 0.55 | 0.44 | 5 | 1 | 357 | `IRSOM` |
| - | - | - | - | - | - | - | - | - | - | - | - | - | - | `SPECtre` |
| ***L. monocytogenes*** **and using overlap of 0.9** | | | | | | | | | | | | | | |
| 1813 | 388 | 191 | 475 | 0.79 | 0.67 | 0.21 | 0.90 | 0.10 | 0.84 | 0.77 | 0 | 0 | 216 | `REPARATION_blast` |
| 36 | 564 | 15 | 2252 | 0.02 | 0.97 | 0.98 | 0.71 | 0.29 | 0.03 | 0.21 | 7 | 2 | 2037 | `Ribo-TISH` |
| 2155 | 224 | 355 | 133 | 0.94 | 0.39 | 0.06 | 0.86 | 0.14 | 0.90 | 0.83 | 0 | 0 | 21984 | `DeepRibo` |
| 975 | 289 | 290 | 1313 | 0.43 | 0.50 | 0.57 | 0.77 | 0.23 | 0.55 | 0.44 | 3 | 1 | 359 | `IRSOM` |
| - | - | - | - | - | - | - | - | - | - | - | - | - | - | `SPECtre` |

**Table S3:** Statistical evaluation of the *translatome* ORFs for the benchmark dataset *P. aeruginosa* and using overlap threshold of 0.01, 0.7, or 0.9. A detailed column description can be found in main Table S1.

| TP | TN | FP | FN | TPR | TNR | FNR | PPV | FDR | F1 | accuracy | sTP | sFP | no_gene | tool |
|---|---|---|---|---|---|---|---|---|---|---|---|---|---|---|
| ***P. aeruginosa*** **and using overlap of 0.01** | | | | | | | | | | | | | | |
| 2315 | 1481 | 154 | 1623 | 0.59 | 0.91 | 0.41 | 0.94 | 0.06 | 0.72 | 0.68 | 49 | 2 | 40 | `REPARATION_blast` |
| 2776 | 1010 | 625 | 1162 | 0.70 | 0.62 | 0.30 | 0.82 | 0.18 | 0.76 | 0.68 | 3143 | 335 | 79 | `Ribo-TISH` |
| 3869 | 1011 | 624 | 69 | 0.98 | 0.62 | 0.02 | 0.86 | 0.14 | 0.92 | 0.88 | 12954 | 790 | 4079 | `DeepRibo` |
| 2813 | 303 | 1332 | 1125 | 0.71 | 0.19 | 0.29 | 0.68 | 0.32 | 0.70 | 0.56 | 322 | 258 | 271 | `IRSOM` |
| 577 | 1178 | 457 | 3361 | 0.15 | 0.72 | 0.85 | 0.56 | 0.44 | 0.23 | 0.31 | 2 | 2 | 0 | `SPECtre` |
| ***P. aeruginosa*** **and using overlap of 0.7** | | | | | | | | | | | | | | |
| 2160 | 1494 | 141 | 1778 | 0.55 | 0.91 | 0.45 | 0.94 | 0.06 | 0.69 | 0.66 | 9 | 0 | 237 | `REPARATION_blast` |
| 143 | 1559 | 76 | 3795 | 0.04 | 0.95 | 0.96 | 0.65 | 0.35 | 0.07 | 0.31 | 79 | 36 | 6615 | `Ribo-TISH` |
| 3689 | 1375 | 260 | 249 | 0.94 | 0.84 | 0.06 | 0.93 | 0.07 | 0.94 | 0.91 | 661 | 45 | 17497 | `DeepRibo` |
| 2608 | 415 | 1220 | 1330 | 0.66 | 0.25 | 0.34 | 0.68 | 0.32 | 0.67 | 0.54 | 14 | 18 | 875 | `IRSOM` |
| 577 | 1178 | 457 | 3361 | 0.15 | 0.72 | 0.85 | 0.56 | 0.44 | 0.23 | 0.31 | 0 | 0 | 0 | `SPECtre` |
| ***P. aeruginosa*** **and using overlap of 0.9** | | | | | | | | | | | | | | |
| 2012 | 1503 | 132 | 1926 | 0.51 | 0.92 | 0.49 | 0.94 | 0.06 | 0.66 | 0.63 | 1 | 0 | 402 | `REPARATION_blast` |
| 71 | 1588 | 47 | 3867 | 0.02 | 0.97 | 0.98 | 0.60 | 0.40 | 0.04 | 0.30 | 12 | 5 | 6814 | `Ribo-TISH` |
| 3571 | 1403 | 232 | 367 | 0.91 | 0.86 | 0.09 | 0.94 | 0.06 | 0.92 | 0.89 | 127 | 15 | 18207 | `DeepRibo` |
| 2607 | 418 | 1217 | 1331 | 0.66 | 0.26 | 0.34 | 0.68 | 0.32 | 0.67 | 0.54 | 6 | 8 | 897 | `IRSOM` |
| 577 | 1178 | 457 | 3361 | 0.15 | 0.72 | 0.85 | 0.56 | 0.44 | 0.23 | 0.31 | 0 | 0 | 0 | `SPECtre` |

**Table S4:** Statistical evaluation of the *translatome* for the benchmark dataset *S.* Typhimurium and using overlap threshold of 0.01, 0.7 or 0.9. A detailed column description can be found in Table S1.

| TP | TN | FP | FN | TPR | TNR | FNR | PPV | FDR | F1 | accuracy | sTP | sFP | no_gene | tool |
|---|---|---|---|---|---|---|---|---|---|---|---|---|---|---|
| ***S.* Typhimurium** and using overlap of **0.01** | | | | | | | | | | | | | | |
| 2998 | 1469 | 212 | 292 | 0.91 | 0.87 | 0.09 | 0.93 | 0.07 | 0.92 | 0.90 | 54 | 8 | 31 | `REPARATION_blast` |
| 2031 | 1446 | 235 | 1259 | 0.62 | 0.86 | 0.38 | 0.90 | 0.10 | 0.73 | 0.70 | 2938 | 162 | 37 | `Ribo-TISH` |
| 3088 | 1456 | 225 | 202 | 0.94 | 0.87 | 0.06 | 0.93 | 0.07 | 0.94 | 0.91 | 15585 | 111 | 1405 | `DeepRibo` |
| 1681 | 852 | 829 | 1609 | 0.51 | 0.51 | 0.49 | 0.67 | 0.33 | 0.58 | 0.51 | 79 | 54 | 88 | `IRSOM` |
| 1017 | 1026 | 655 | 2275 | 0.31 | 0.61 | 0.69 | 0.61 | 0.39 | 0.41 | 0.41 | 15 | 16 | 0 | `SPECtre` |
| ***S.* Typhimurium** and using overlap of **0.7** | | | | | | | | | | | | | | |
| 2961 | 1544 | 137 | 329 | 0.90 | 0.92 | 0.10 | 0.96 | 0.04 | 0.93 | 0.91 | 0 | 0 | 127 | `REPARATION_blast` |
| 323 | 1594 | 87 | 2967 | 0.10 | 0.95 | 0.90 | 0.79 | 0.21 | 0.17 | 0.39 | 256 | 40 | 4669 | `Ribo-TISH` |
| 2522 | 1651 | 30 | 768 | 0.77 | 0.98 | 0.23 | 0.99 | 0.01 | 0.86 | 0.84 | 19 | 1 | 17738 | `DeepRibo` |
| 1647 | 896 | 785 | 1643 | 0.50 | 0.53 | 0.50 | 0.68 | 0.32 | 0.58 | 0.51 | 10 | 9 | 182 | `IRSOM` |
| 1017 | 1042 | 639 | 2275 | 0.31 | 0.62 | 0.69 | 0.61 | 0.39 | 0.41 | 0.41 | 0 | 0 | 0 | `SPECtre` |
| ***S.* Typhimurium** and using overlap of **0.9** | | | | | | | | | | | | | | |
| 2859 | 1556 | 125 | 431 | 0.87 | 0.93 | 0.13 | 0.96 | 0.04 | 0.91 | 0.89 | 0 | 0 | 241 | `REPARATION_blast` |
| 221 | 1621 | 60 | 3069 | 0.07 | 0.96 | 0.93 | 0.79 | 0.21 | 0.12 | 0.37 | 42 | 8 | 5044 | `Ribo-TISH` |
| 2459 | 1655 | 26 | 831 | 0.75 | 0.98 | 0.25 | 0.99 | 0.01 | 0.85 | 0.83 | 1 | 0 | 17824 | `DeepRibo` |
| 1646 | 896 | 785 | 1644 | 0.50 | 0.53 | 0.50 | 0.68 | 0.32 | 0.58 | 0.51 | 3 | 1 | 198 | `IRSOM` |
| 1017 | 1042 | 639 | 2275 | 0.31 | 0.62 | 0.69 | 0.61 | 0.39 | 0.41 | 0.41 | 0 | 0 | 0 | `SPECtre` |

#### D.2 Operons

**Table S5:** Statistical evaluation of ORFs located in operons for the benchmark dataset *E. coli* and using overlap threshold of 0.01, 0.7, or 0.9. A detailed column description can be found in main Table S1.

| TP | TN | FP | FN | TPR | TNR | FNR | PPV | FDR | F1 | accuracy | sTP | sFP | tool |
|---|---|---|---|---|---|---|---|---|---|---|---|---|---|
| ***E. coli*** **and using overlap of 0.01** | | | | | | | | | | | | | |
| 1767 | 448 | 566 | 28 | 0.98 | 0.44 | 0.02 | 0.76 | 0.24 | 0.86 | 0.79 | 37 | 8 | REPARATION_blast |
| 1127 | 839 | 175 | 668 | 0.63 | 0.83 | 0.37 | 0.87 | 0.13 | 0.73 | 0.70 | 1666 | 97 | Ribo-TISH |
| 1727 | 848 | 166 | 68 | 0.96 | 0.84 | 0.04 | 0.91 | 0.09 | 0.94 | 0.92 | 10444 | 164 | DeepRibo |
| 902 | 515 | 499 | 893 | 0.50 | 0.51 | 0.50 | 0.64 | 0.36 | 0.56 | 0.50 | 21 | 16 | IRSOM |
| 698 | 548 | 466 | 1097 | 0.39 | 0.54 | 0.61 | 0.60 | 0.40 | 0.47 | 0.44 | 0 | 0 | SPECtre |
| ***E. coli*** **and using overlap of 0.7** | | | | | | | | | | | | | |
| 1757 | 493 | 521 | 38 | 0.98 | 0.49 | 0.02 | 0.77 | 0.23 | 0.86 | 0.80 | 0 | 0 | REPARATION_blast |
| 54 | 968 | 46 | 1741 | 0.03 | 0.95 | 0.97 | 0.54 | 0.46 | 0.06 | 0.36 | 38 | 12 | Ribo-TISH |
| 1490 | 985 | 29 | 305 | 0.83 | 0.97 | 0.17 | 0.98 | 0.02 | 0.90 | 0.88 | 8 | 0 | DeepRibo |
| 892 | 527 | 487 | 903 | 0.50 | 0.52 | 0.50 | 0.65 | 0.35 | 0.56 | 0.51 | 1 | 4 | IRSOM |
| 697 | 552 | 462 | 1098 | 0.39 | 0.54 | 0.61 | 0.60 | 0.40 | 0.47 | 0.44 | 0 | 0 | SPECtre |
| ***E. coli*** **and using overlap of 0.9** | | | | | | | | | | | | | |
| 1726 | 529 | 485 | 69 | 0.96 | 0.52 | 0.04 | 0.78 | 0.22 | 0.86 | 0.80 | 0 | 0 | REPARATION_blast |
| 37 | 989 | 25 | 1758 | 0.02 | 0.98 | 0.98 | 0.60 | 0.40 | 0.04 | 0.37 | 7 | 2 | Ribo-TISH |
| 1456 | 987 | 27 | 339 | 0.81 | 0.97 | 0.19 | 0.98 | 0.02 | 0.89 | 0.87 | 1 | 0 | DeepRibo |
| 892 | 527 | 487 | 903 | 0.50 | 0.52 | 0.50 | 0.65 | 0.35 | 0.56 | 0.51 | 1 | 1 | IRSOM |
| 697 | 552 | 462 | 1098 | 0.39 | 0.54 | 0.61 | 0.60 | 0.40 | 0.47 | 0.44 | 0 | 0 | SPECtre |

**Table S6:** Statistical evaluation of ORFs located in operons for the benchmark dataset *L. monocytogenes* and using overlap threshold of 0.01, 0.7 or 0.9. A detailed column description can be found in Table S1.

| TP | TN | FP | FN | TPR | TNR | FNR | PPV | FDR | F1 | accuracy | sTP | sFP | tool |
|---|---|---|---|---|---|---|---|---|---|---|---|---|---|
| ***L. monocytogenes*** **and using overlap of 0.01** | | | | | | | | | | | | | |
| 1374 | 253 | 179 | 248 | 0.85 | 0.59 | 0.15 | 0.88 | 0.12 | 0.87 | 0.79 | 34 | 4 | REPARATION_blast |
| 723 | 291 | 141 | 899 | 0.45 | 0.67 | 0.55 | 0.84 | 0.16 | 0.58 | 0.49 | 472 | 82 | Ribo-TISH |
| 1619 | 45 | 387 | 3 | 1.00 | 0.10 | 0.00 | 0.81 | 0.19 | 0.89 | 0.81 | 10842 | 1537 | DeepRibo |
| 721 | 187 | 245 | 901 | 0.44 | 0.43 | 0.56 | 0.75 | 0.25 | 0.56 | 0.44 | 59 | 37 | IRSOM |
| - | - | - | - | - | - | - | - | - | - | - | - | - | SPECtre |
| ***L. monocytogenes*** **and using overlap of 0.7** | | | | | | | | | | | | | |
| 1322 | 274 | 158 | 300 | 0.82 | 0.63 | 0.18 | 0.89 | 0.11 | 0.85 | 0.78 | 0 | 0 | REPARATION_blast |
| 36 | 416 | 16 | 1586 | 0.02 | 0.96 | 0.98 | 0.69 | 0.31 | 0.04 | 0.22 | 21 | 12 | Ribo-TISH |
| 1554 | 159 | 273 | 68 | 0.96 | 0.37 | 0.04 | 0.85 | 0.15 | 0.90 | 0.83 | 2 | 0 | DeepRibo |
| 665 | 208 | 224 | 957 | 0.41 | 0.48 | 0.59 | 0.75 | 0.25 | 0.53 | 0.43 | 1 | 0 | IRSOM |
| - | - | - | - | - | - | - | - | - | - | - | - | - | SPECtre |
| ***L. monocytogenes*** **and using overlap of 0.9** | | | | | | | | | | | | | |
| 1300 | 282 | 150 | 322 | 0.80 | 0.65 | 0.20 | 0.90 | 0.10 | 0.85 | 0.77 | 0 | 0 | REPARATION_blast |
| 21 | 421 | 11 | 1601 | 0.01 | 0.97 | 0.99 | 0.66 | 0.34 | 0.03 | 0.22 | 6 | 2 | Ribo-TISH |
| 1526 | 164 | 268 | 96 | 0.94 | 0.38 | 0.06 | 0.85 | 0.15 | 0.89 | 0.82 | 0 | 0 | DeepRibo |
| 665 | 208 | 224 | 957 | 0.41 | 0.48 | 0.59 | 0.75 | 0.25 | 0.53 | 0.43 | 0 | 0 | IRSOM |
| - | - | - | - | - | - | - | - | - | - | - | - | - | SPECtre |

**Table S7:** Statistical evaluation of ORFs located in operons for the benchmark dataset *P. aeruginosa* and using overlap threshold of 0.01, 0.7 or 0.9. A detailed column description can be found in Table S1.

| TP | TN | FP | FN | TPR | TNR | FNR | PPV | FDR | F1 | accuracy | sTP | sFP | tool |
|---|---|---|---|---|---|---|---|---|---|---|---|---|---|
| ***P. aeruginosa*** **and using overlap of 0.01** | | | | | | | | | | | | | |
| 1586 | 1009 | 103 | 926 | 0.63 | 0.91 | 0.37 | 0.94 | 0.06 | 0.76 | 0.72 | 29 | 2 | REPARATION_blast |
| 1819 | 699 | 413 | 693 | 0.72 | 0.63 | 0.28 | 0.81 | 0.19 | 0.77 | 0.69 | 2196 | 234 | Ribo-TISH |
| 2473 | 715 | 397 | 39 | 0.98 | 0.64 | 0.02 | 0.86 | 0.14 | 0.92 | 0.88 | 8365 | 510 | DeepRibo |
| 1760 | 204 | 908 | 752 | 0.70 | 0.18 | 0.30 | 0.66 | 0.34 | 0.68 | 0.54 | 178 | 159 | IRSOM |
| 340 | 749 | 363 | 2172 | 0.14 | 0.67 | 0.86 | 0.48 | 0.52 | 0.21 | 0.30 | 2 | 2 | SPECtre |
| ***P. aeruginosa*** **and using overlap of 0.7** | | | | | | | | | | | | | |
| 1494 | 1018 | 94 | 1018 | 0.59 | 0.92 | 0.41 | 0.94 | 0.06 | 0.73 | 0.69 | 6 | 0 | REPARATION_blast |
| 81 | 1061 | 51 | 2431 | 0.03 | 0.95 | 0.97 | 0.61 | 0.39 | 0.06 | 0.32 | 43 | 30 | Ribo-TISH |
| 2353 | 947 | 165 | 159 | 0.94 | 0.85 | 0.06 | 0.93 | 0.07 | 0.94 | 0.91 | 361 | 25 | DeepRibo |
| 1644 | 270 | 842 | 868 | 0.65 | 0.24 | 0.35 | 0.66 | 0.34 | 0.66 | 0.53 | 4 | 7 | IRSOM |
| 340 | 750 | 362 | 2172 | 0.14 | 0.67 | 0.86 | 0.48 | 0.52 | 0.21 | 0.30 | 0 | 0 | SPECtre |
| ***P. aeruginosa*** **and using overlap of 0.9** | | | | | | | | | | | | | |
| 1401 | 1025 | 87 | 1111 | 0.56 | 0.92 | 0.44 | 0.94 | 0.06 | 0.70 | 0.67 | 1 | 0 | REPARATION_blast |
| 41 | 1079 | 33 | 2471 | 0.02 | 0.97 | 0.98 | 0.55 | 0.45 | 0.03 | 0.31 | 9 | 5 | Ribo-TISH |
| 2298 | 964 | 148 | 214 | 0.91 | 0.87 | 0.09 | 0.94 | 0.06 | 0.93 | 0.90 | 81 | 7 | DeepRibo |
| 1643 | 271 | 841 | 869 | 0.65 | 0.24 | 0.35 | 0.66 | 0.34 | 0.66 | 0.53 | 0 | 4 | IRSOM |
| 340 | 750 | 362 | 2172 | 0.14 | 0.67 | 0.86 | 0.48 | 0.52 | 0.21 | 0.30 | 0 | 0 | SPECtre |

**Table S8:** Statistical evaluation of ORFs located in operons for the benchmark dataset *S.* Typhimurium and using overlap threshold of 0.01, 0.7 or 0.9. A detailed column description can be found in Table S1.

| TP | TN | FP | FN | TPR | TNR | FNR | PPV | FDR | F1 | accuracy | sTP | sFP | tool |
|---|---|---|---|---|---|---|---|---|---|---|---|---|---|
| ***S.* Typhimurium** and using overlap of **0.01** | | | | | | | | | | | | | |
| 1819 | 861 | 141 | 134 | 0.93 | 0.86 | 0.07 | 0.93 | 0.07 | 0.93 | 0.91 | 41 | 8 | `REPARATION_blast` |
| 1235 | 836 | 166 | 718 | 0.63 | 0.83 | 0.37 | 0.88 | 0.12 | 0.74 | 0.70 | 1784 | 108 | `Ribo-TISH` |
| 1833 | 876 | 126 | 120 | 0.94 | 0.87 | 0.06 | 0.94 | 0.06 | 0.94 | 0.92 | 9536 | 57 | `DeepRibo` |
| 983 | 461 | 541 | 970 | 0.50 | 0.46 | 0.50 | 0.65 | 0.35 | 0.57 | 0.49 | 41 | 44 | `IRSOM` |
| 555 | 554 | 448 | 1398 | 0.28 | 0.55 | 0.72 | 0.55 | 0.45 | 0.38 | 0.38 | 12 | 11 | `SPECtre` |
| ***S.* Typhimurium** and using overlap of **0.7** | | | | | | | | | | | | | |
| 1796 | 901 | 101 | 157 | 0.92 | 0.90 | 0.08 | 0.95 | 0.05 | 0.93 | 0.91 | 0 | 0 | `REPARATION_blast` |
| 178 | 944 | 58 | 1775 | 0.09 | 0.94 | 0.91 | 0.75 | 0.25 | 0.16 | 0.38 | 139 | 28 | `Ribo-TISH` |
| 1482 | 991 | 11 | 471 | 0.76 | 0.99 | 0.24 | 0.99 | 0.01 | 0.86 | 0.84 | 7 | 0 | `DeepRibo` |
| 969 | 469 | 533 | 984 | 0.50 | 0.47 | 0.50 | 0.65 | 0.35 | 0.56 | 0.49 | 2 | 6 | `IRSOM` |
| 554 | 561 | 441 | 1399 | 0.28 | 0.56 | 0.72 | 0.56 | 0.44 | 0.38 | 0.38 | 0 | 0 | `SPECtre` |
| ***S.* Typhimurium** and using overlap of **0.9** | | | | | | | | | | | | | |
| 1746 | 912 | 90 | 207 | 0.89 | 0.91 | 0.11 | 0.95 | 0.05 | 0.92 | 0.90 | 0 | 0 | `REPARATION_blast` |
| 123 | 963 | 39 | 1830 | 0.06 | 0.96 | 0.94 | 0.76 | 0.24 | 0.12 | 0.37 | 24 | 7 | `Ribo-TISH` |
| 1459 | 992 | 10 | 494 | 0.75 | 0.99 | 0.25 | 0.99 | 0.01 | 0.85 | 0.83 | 1 | 0 | `DeepRibo` |
| 968 | 469 | 533 | 985 | 0.50 | 0.47 | 0.50 | 0.64 | 0.36 | 0.56 | 0.49 | 1 | 1 | `IRSOM` |
| 554 | 561 | 441 | 1399 | 0.28 | 0.56 | 0.72 | 0.56 | 0.44 | 0.38 | 0.38 | 0 | 0 | `SPECtre` |

##### D.3 Operons complement

**Table S9:** Statistical evaluation of ORFs located outside operons for the benchmark dataset *E. coli* and using overlap threshold of 0.01, 0.7, or 0.9. A detailed column description can be found in main Table S1.

| TP | TN | FP | FN | TPR | TNR | FNR | PPV | FDR | F1 | accuracy | sTP | sFP | tool |
|---|---|---|---|---|---|---|---|---|---|---|---|---|---|
| ***E. coli*** **and using overlap of 0.01** | | | | | | | | | | | | | |
| 944 | 190 | 279 | 26 | 0.97 | 0.41 | 0.03 | 0.77 | 0.23 | 0.86 | 0.79 | 20 | 3 | REPARATION_blast |
| 624 | 375 | 94 | 346 | 0.64 | 0.80 | 0.36 | 0.87 | 0.13 | 0.74 | 0.69 | 907 | 60 | Ribo-TISH |
| 943 | 355 | 114 | 27 | 0.97 | 0.76 | 0.03 | 0.89 | 0.11 | 0.93 | 0.90 | 5892 | 178 | DeepRibo |
| 552 | 240 | 229 | 418 | 0.57 | 0.51 | 0.43 | 0.71 | 0.29 | 0.63 | 0.55 | 34 | 19 | IRSOM |
| 390 | 259 | 210 | 580 | 0.40 | 0.55 | 0.60 | 0.65 | 0.35 | 0.50 | 0.45 | 0 | 0 | SPECtre |
| ***E. coli*** **and using overlap of 0.7** | | | | | | | | | | | | | |
| 941 | 219 | 250 | 29 | 0.97 | 0.47 | 0.03 | 0.79 | 0.21 | 0.87 | 0.81 | 1 | 0 | REPARATION_blast |
| 30 | 446 | 23 | 940 | 0.03 | 0.95 | 0.97 | 0.57 | 0.43 | 0.06 | 0.33 | 21 | 14 | Ribo-TISH |
| 816 | 450 | 19 | 154 | 0.84 | 0.96 | 0.16 | 0.98 | 0.02 | 0.90 | 0.88 | 9 | 1 | DeepRibo |
| 516 | 255 | 214 | 454 | 0.53 | 0.54 | 0.47 | 0.71 | 0.29 | 0.61 | 0.54 | 5 | 3 | IRSOM |
| 390 | 260 | 209 | 580 | 0.40 | 0.55 | 0.60 | 0.65 | 0.35 | 0.50 | 0.45 | 0 | 0 | SPECtre |
| ***E. coli*** **and using overlap of 0.9** | | | | | | | | | | | | | |
| 930 | 237 | 232 | 40 | 0.96 | 0.51 | 0.04 | 0.80 | 0.20 | 0.87 | 0.81 | 0 | 0 | REPARATION_blast |
| 14 | 453 | 16 | 956 | 0.01 | 0.97 | 0.99 | 0.47 | 0.53 | 0.03 | 0.32 | 2 | 2 | Ribo-TISH |
| 790 | 451 | 18 | 180 | 0.81 | 0.96 | 0.19 | 0.98 | 0.02 | 0.89 | 0.86 | 2 | 0 | DeepRibo |
| 516 | 255 | 214 | 454 | 0.53 | 0.54 | 0.47 | 0.71 | 0.29 | 0.61 | 0.54 | 1 | 1 | IRSOM |
| 390 | 260 | 209 | 580 | 0.40 | 0.55 | 0.60 | 0.65 | 0.35 | 0.50 | 0.45 | 0 | 0 | SPECtre |

**Table S10:** Statistical evaluation of ORFs located outside operons for the benchmark dataset *L. monocytogenes* and using overlap threshold of 0.01, 0.7 or 0.9. A detailed column description can be found in Table S1.

| TP | TN | FP | FN | TPR | TNR | FNR | PPV | FDR | F1 | accuracy | sTP | sFP | tool |
|---|---|---|---|---|---|---|---|---|---|---|---|---|---|
| ***L. monocytogenes*** **and using overlap of 0.01** | | | | | | | | | | | | | |
| 540 | 96 | 51 | 126 | 0.81 | 0.65 | 0.19 | 0.91 | 0.09 | 0.86 | 0.78 | 14 | 0 | REPARATION_blast |
| 300 | 99 | 48 | 366 | 0.45 | 0.67 | 0.55 | 0.86 | 0.14 | 0.59 | 0.49 | 223 | 28 | Ribo-TISH |
| 665 | 11 | 136 | 1 | 1.00 | 0.07 | 0.00 | 0.83 | 0.17 | 0.91 | 0.83 | 4595 | 514 | DeepRibo |
| 395 | 57 | 90 | 271 | 0.59 | 0.39 | 0.41 | 0.81 | 0.19 | 0.69 | 0.56 | 81 | 18 | IRSOM |
| - | - | - | - | - | - | - | - | - | - | - | - | - | SPECtre |
| ***L. monocytogenes*** **and using overlap of 0.7** | | | | | | | | | | | | | |
| 521 | 104 | 43 | 145 | 0.78 | 0.71 | 0.22 | 0.92 | 0.08 | 0.85 | 0.77 | 0 | 0 | REPARATION_blast |
| 19 | 140 | 7 | 647 | 0.03 | 0.95 | 0.97 | 0.73 | 0.27 | 0.05 | 0.20 | 11 | 6 | Ribo-TISH |
| 640 | 56 | 91 | 26 | 0.96 | 0.38 | 0.04 | 0.88 | 0.12 | 0.92 | 0.86 | 2 | 1 | DeepRibo |
| 310 | 81 | 66 | 356 | 0.47 | 0.55 | 0.53 | 0.82 | 0.18 | 0.60 | 0.48 | 4 | 1 | IRSOM |
| - | - | - | - | - | - | - | - | - | - | - | - | - | SPECtre |
| ***L. monocytogenes*** **and using overlap of 0.9** | | | | | | | | | | | | | |
| 513 | 106 | 41 | 153 | 0.77 | 0.72 | 0.23 | 0.93 | 0.07 | 0.84 | 0.76 | 0 | 0 | REPARATION_blast |
| 15 | 143 | 4 | 651 | 0.02 | 0.97 | 0.98 | 0.79 | 0.21 | 0.04 | 0.19 | 1 | 0 | Ribo-TISH |
| 629 | 60 | 87 | 37 | 0.94 | 0.41 | 0.06 | 0.88 | 0.12 | 0.91 | 0.85 | 0 | 0 | DeepRibo |
| 310 | 81 | 66 | 356 | 0.47 | 0.55 | 0.53 | 0.82 | 0.18 | 0.60 | 0.48 | 3 | 1 | IRSOM |
| - | - | - | - | - | - | - | - | - | - | - | - | - | SPECtre |

**Table S11:** Statistical evaluation of ORFs located outside operons for the benchmark dataset *P. aeruginosa* and using overlap threshold of 0.01, 0.7, or 0.9. A detailed column description can be found in main Table S1.

| TP | TN | FP | FN | TPR | TNR | FNR | PPV | FDR | F1 | accuracy | sTP | sFP | tool |
|---|---|---|---|---|---|---|---|---|---|---|---|---|---|
| ***P. aeruginosa*** **and using overlap of 0.01** | | | | | | | | | | | | | |
| 730 | 472 | 51 | 696 | 0.51 | 0.90 | 0.49 | 0.93 | 0.07 | 0.66 | 0.62 | 27 | 0 | REPARATION_blast |
| 957 | 311 | 212 | 469 | 0.67 | 0.59 | 0.33 | 0.82 | 0.18 | 0.74 | 0.65 | 947 | 101 | Ribo-TISH |
| 1396 | 296 | 227 | 30 | 0.98 | 0.57 | 0.02 | 0.86 | 0.14 | 0.92 | 0.87 | 4650 | 280 | DeepRibo |
| 1095 | 95 | 428 | 331 | 0.77 | 0.18 | 0.23 | 0.72 | 0.28 | 0.74 | 0.61 | 250 | 145 | IRSOM |
| 237 | 428 | 95 | 1189 | 0.17 | 0.82 | 0.83 | 0.71 | 0.29 | 0.27 | 0.34 | 0 | 0 | SPECtre |
| ***P. aeruginosa*** **and using overlap of 0.7** | | | | | | | | | | | | | |
| 666 | 476 | 47 | 760 | 0.47 | 0.91 | 0.53 | 0.93 | 0.07 | 0.62 | 0.59 | 3 | 0 | REPARATION_blast |
| 62 | 498 | 25 | 1364 | 0.04 | 0.95 | 0.96 | 0.71 | 0.29 | 0.08 | 0.29 | 36 | 6 | Ribo-TISH |
| 1336 | 428 | 95 | 90 | 0.94 | 0.82 | 0.06 | 0.93 | 0.07 | 0.94 | 0.91 | 300 | 20 | DeepRibo |
| 964 | 145 | 378 | 462 | 0.68 | 0.28 | 0.32 | 0.72 | 0.28 | 0.70 | 0.57 | 10 | 11 | IRSOM |
| 237 | 428 | 95 | 1189 | 0.17 | 0.82 | 0.83 | 0.71 | 0.29 | 0.27 | 0.34 | 0 | 0 | SPECtre |
| ***P. aeruginosa*** **and using overlap of 0.9** | | | | | | | | | | | | | |
| 611 | 478 | 45 | 815 | 0.43 | 0.91 | 0.57 | 0.93 | 0.07 | 0.59 | 0.56 | 0 | 0 | REPARATION_blast |
| 30 | 509 | 14 | 1396 | 0.02 | 0.97 | 0.98 | 0.68 | 0.32 | 0.04 | 0.28 | 3 | 0 | Ribo-TISH |
| 1273 | 439 | 84 | 153 | 0.89 | 0.84 | 0.11 | 0.94 | 0.06 | 0.91 | 0.88 | 46 | 8 | DeepRibo |
| 964 | 147 | 376 | 462 | 0.68 | 0.28 | 0.32 | 0.72 | 0.28 | 0.70 | 0.57 | 6 | 4 | IRSOM |
| 237 | 428 | 95 | 1189 | 0.17 | 0.82 | 0.83 | 0.71 | 0.29 | 0.27 | 0.34 | 0 | 0 | SPECtre |

**Table S12:** Statistical evaluation of ORFs located outside operons for the benchmark dataset *S.* Typhimurium and using overlap threshold of 0.01, 0.7, or 0.9. A detailed column description can be found in Table S1.

| TP | TN | FP | FN | TPR | TNR | FNR | PPV | FDR | F1 | accuracy | sTP | sFP | tool |
|---|---|---|---|---|---|---|---|---|---|---|---|---|---|
| ***S.* Typhimurium** and using overlap of **0.01** | | | | | | | | | | | | | |
| 1184 | 607 | 72 | 153 | 0.89 | 0.89 | 0.11 | 0.94 | 0.06 | 0.91 | 0.89 | 24 | 0 | `REPARATION_blast` |
| 796 | 609 | 70 | 541 | 0.60 | 0.90 | 0.40 | 0.92 | 0.08 | 0.72 | 0.70 | 1166 | 54 | `Ribo-TISH` |
| 1257 | 580 | 99 | 80 | 0.94 | 0.85 | 0.06 | 0.93 | 0.07 | 0.93 | 0.91 | 6079 | 55 | `DeepRibo` |
| 702 | 389 | 290 | 635 | 0.53 | 0.57 | 0.47 | 0.71 | 0.29 | 0.60 | 0.54 | 47 | 18 | `IRSOM` |
| 463 | 470 | 209 | 874 | 0.35 | 0.69 | 0.65 | 0.69 | 0.31 | 0.46 | 0.46 | 5 | 5 | `SPECtre` |
| ***S.* Typhimurium** and using overlap of **0.7** | | | | | | | | | | | | | |
| 1165 | 643 | 36 | 172 | 0.87 | 0.95 | 0.13 | 0.97 | 0.03 | 0.92 | 0.90 | 0 | 0 | `REPARATION_blast` |
| 145 | 650 | 29 | 1192 | 0.11 | 0.96 | 0.89 | 0.83 | 0.17 | 0.19 | 0.39 | 117 | 12 | `Ribo-TISH` |
| 1040 | 660 | 19 | 297 | 0.78 | 0.97 | 0.22 | 0.98 | 0.02 | 0.87 | 0.84 | 12 | 1 | `DeepRibo` |
| 678 | 427 | 252 | 659 | 0.51 | 0.63 | 0.49 | 0.73 | 0.27 | 0.60 | 0.55 | 8 | 3 | `IRSOM` |
| 461 | 481 | 198 | 876 | 0.34 | 0.71 | 0.66 | 0.70 | 0.30 | 0.46 | 0.47 | 0 | 0 | `SPECtre` |
| ***S.* Typhimurium** and using overlap of **0.9** | | | | | | | | | | | | | |
| 1113 | 644 | 35 | 224 | 0.83 | 0.95 | 0.17 | 0.97 | 0.03 | 0.90 | 0.87 | 0 | 0 | `REPARATION_blast` |
| 98 | 658 | 21 | 1239 | 0.07 | 0.97 | 0.93 | 0.82 | 0.18 | 0.13 | 0.38 | 18 | 1 | `Ribo-TISH` |
| 1000 | 663 | 16 | 337 | 0.75 | 0.98 | 0.25 | 0.98 | 0.02 | 0.85 | 0.82 | 0 | 0 | `DeepRibo` |
| 678 | 427 | 252 | 659 | 0.51 | 0.63 | 0.49 | 0.73 | 0.27 | 0.60 | 0.55 | 2 | 0 | `IRSOM` |
| 461 | 481 | 198 | 876 | 0.34 | 0.71 | 0.66 | 0.70 | 0.30 | 0.46 | 0.47 | 0 | 0 | `SPECtre` |

#### D.4 Small Open Reading Frames

**Table S13:** Statistical evaluation of sORFs for the benchmark dataset *E. coli*, *P. aeruginosa* and *S.* Typhimurium and using overlap threshold of 0.01, 0.7 or 0.9. We could not find any sORF results for the benchmark dataset: *L. monocytogenes*. A detailed column description can be found in Table S1.

| TP | TN | FP | FN | TPR | TNR | FNR | PPV | FDR | F1 | accuracy | sTP | sFP | tool |
|---|---|---|---|---|---|---|---|---|---|---|---|---|---|
| ***E. coli*** **and using overlap of 0.01** | | | | | | | | | | | | | |
| 25 | 51 | 8 | 30 | 0.45 | 0.86 | 0.55 | 0.76 | 0.24 | 0.57 | 0.67 | 0 | 0 | `REPARATION_blast` |
| 7 | 58 | 1 | 48 | 0.13 | 0.98 | 0.87 | 0.88 | 0.12 | 0.22 | 0.57 | 0 | 0 | `Ribo-TISH` |
| 50 | 49 | 10 | 5 | 0.91 | 0.83 | 0.09 | 0.83 | 0.17 | 0.87 | 0.87 | 55 | 8 | `DeepRibo` |
| 9 | 47 | 12 | 46 | 0.16 | 0.80 | 0.84 | 0.43 | 0.57 | 0.24 | 0.49 | 0 | 1 | `IRSOM` |
| 21 | 35 | 24 | 34 | 0.38 | 0.59 | 0.62 | 0.47 | 0.53 | 0.42 | 0.49 | 0 | 1 | `SPECtre` |
| ***E. coli*** **and using overlap of 0.7** | | | | | | | | | | | | | |
| 19 | 53 | 6 | 36 | 0.35 | 0.90 | 0.65 | 0.76 | 0.24 | 0.48 | 0.63 | 0 | 0 | `REPARATION_blast` |
| 3 | 59 | 0 | 52 | 0.05 | 1.00 | 0.95 | 1.00 | 0.00 | 0.10 | 0.54 | 0 | 0 | `Ribo-TISH` |
| 44 | 52 | 7 | 11 | 0.80 | 0.88 | 0.20 | 0.86 | 0.14 | 0.83 | 0.84 | 3 | 0 | `DeepRibo` |
| 4 | 50 | 9 | 51 | 0.07 | 0.85 | 0.93 | 0.31 | 0.69 | 0.12 | 0.47 | 0 | 0 | `IRSOM` |
| 19 | 35 | 24 | 36 | 0.35 | 0.59 | 0.65 | 0.44 | 0.56 | 0.39 | 0.47 | 0 | 0 | `SPECtre` |
| ***E. coli*** **and using overlap of 0.9** | | | | | | | | | | | | | |
| 19 | 53 | 6 | 36 | 0.35 | 0.90 | 0.65 | 0.76 | 0.24 | 0.48 | 0.63 | 0 | 0 | `REPARATION_blast` |
| 2 | 59 | 0 | 53 | 0.04 | 1.00 | 0.96 | 1.00 | 0.00 | 0.07 | 0.54 | 0 | 0 | `Ribo-TISH` |
| 42 | 52 | 7 | 13 | 0.76 | 0.88 | 0.24 | 0.86 | 0.14 | 0.81 | 0.82 | 2 | 0 | `DeepRibo` |
| 4 | 50 | 9 | 51 | 0.07 | 0.85 | 0.93 | 0.31 | 0.69 | 0.12 | 0.47 | 0 | 0 | `IRSOM` |
| 19 | 35 | 24 | 36 | 0.35 | 0.59 | 0.65 | 0.44 | 0.56 | 0.39 | 0.47 | 0 | 0 | `SPECtre` |
| ***P. aeruginosa*** **and using overlap of 0.01** | | | | | | | | | | | | | |
| 5 | 5 | 0 | 2 | 0.71 | 1.0 | 0.29 | 1.00 | 0.00 | 0.83 | 0.83 | 0 | 0 | `REPARATION_blast` |
| 1 | 5 | 0 | 6 | 0.14 | 1.0 | 0.86 | 1.00 | 0.00 | 0.25 | 0.50 | 0 | 0 | `Ribo-TISH` |
| 7 | 4 | 1 | 0 | 1.00 | 0.8 | 0.00 | 0.88 | 0.12 | 0.93 | 0.92 | 6 | 2 | `DeepRibo` |
| 4 | 4 | 1 | 3 | 0.57 | 0.8 | 0.43 | 0.80 | 0.20 | 0.67 | 0.67 | 0 | 0 | `IRSOM` |
| 1 | 2 | 3 | 6 | 0.14 | 0.4 | 0.86 | 0.25 | 0.75 | 0.18 | 0.25 | 0 | 0 | `SPECtre` |
| ***P. aeruginosa*** **and using overlap of 0.7** | | | | | | | | | | | | | |
| 4 | 5 | 0 | 3 | 0.57 | 1.0 | 0.43 | 1.00 | 0.00 | 0.73 | 0.75 | 0 | 0 | `REPARATION_blast` |
| 0 | 5 | 0 | 7 | 0.00 | 1.0 | 1.00 | 0.00 | 0.00 | 0.00 | 0.42 | 0 | 0 | `Ribo-TISH` |
| 7 | 4 | 1 | 0 | 1.00 | 0.8 | 0.00 | 0.88 | 0.12 | 0.93 | 0.92 | 0 | 1 | `DeepRibo` |
| 3 | 5 | 0 | 4 | 0.43 | 1.0 | 0.57 | 1.00 | 0.00 | 0.60 | 0.67 | 0 | 0 | `IRSOM` |
| 1 | 2 | 3 | 6 | 0.14 | 0.4 | 0.86 | 0.25 | 0.75 | 0.18 | 0.25 | 0 | 0 | `SPECtre` |
| ***P. aeruginosa*** **and using overlap of 0.9** | | | | | | | | | | | | | |
| 4 | 5 | 0 | 3 | 0.57 | 1.0 | 0.43 | 1.00 | 0.00 | 0.73 | 0.75 | 0 | 0 | `REPARATION_blast` |
| 0 | 5 | 0 | 7 | 0.00 | 1.0 | 1.00 | 0.00 | 0.00 | 0.00 | 0.42 | 0 | 0 | `Ribo-TISH` |
| 6 | 4 | 1 | 1 | 0.86 | 0.8 | 0.14 | 0.86 | 0.14 | 0.86 | 0.83 | 0 | 0 | `DeepRibo` |
| 3 | 5 | 0 | 4 | 0.43 | 1.0 | 0.57 | 1.00 | 0.00 | 0.60 | 0.67 | 0 | 0 | `IRSOM` |
| 1 | 2 | 3 | 6 | 0.14 | 0.4 | 0.86 | 0.25 | 0.75 | 0.18 | 0.25 | 0 | 0 | `SPECtre` |
| ***S.*** **Typhimurium and using overlap of 0.01** | | | | | | | | | | | | | |
| 10 | 61 | 8 | 21 | 0.32 | 0.88 | 0.68 | 0.56 | 0.44 | 0.41 | 0.71 | 0 | 0 | `REPARATION_blast` |
| 4 | 64 | 5 | 27 | 0.13 | 0.93 | 0.87 | 0.44 | 0.56 | 0.20 | 0.68 | 0 | 0 | `Ribo-TISH` |
| 30 | 59 | 10 | 1 | 0.97 | 0.86 | 0.03 | 0.75 | 0.25 | 0.85 | 0.89 | 38 | 11 | `DeepRibo` |
| 6 | 48 | 21 | 25 | 0.19 | 0.70 | 0.81 | 0.22 | 0.78 | 0.21 | 0.54 | 1 | 1 | `IRSOM` |
| 10 | 44 | 25 | 21 | 0.32 | 0.64 | 0.68 | 0.29 | 0.71 | 0.30 | 0.54 | 0 | 2 | `SPECtre` |
| ***S.*** **Typhimurium and using overlap of 0.7** | | | | | | | | | | | | | |
| 9 | 67 | 2 | 22 | 0.29 | 0.97 | 0.71 | 0.82 | 0.18 | 0.43 | 0.76 | 0 | 0 | `REPARATION_blast` |
| 2 | 68 | 1 | 29 | 0.06 | 0.99 | 0.94 | 0.67 | 0.33 | 0.12 | 0.70 | 0 | 0 | `Ribo-TISH` |
| 26 | 64 | 5 | 5 | 0.84 | 0.93 | 0.16 | 0.84 | 0.16 | 0.84 | 0.90 | 1 | 0 | `DeepRibo` |
| 5 | 58 | 11 | 26 | 0.16 | 0.84 | 0.84 | 0.31 | 0.69 | 0.21 | 0.63 | 0 | 0 | `IRSOM` |
| 9 | 48 | 21 | 22 | 0.29 | 0.70 | 0.71 | 0.30 | 0.70 | 0.30 | 0.57 | 0 | 0 | `SPECtre` |
| ***S.*** **Typhimurium and using overlap of 0.9** | | | | | | | | | | | | | |
| 9 | 68 | 1 | 22 | 0.29 | 0.99 | 0.71 | 0.90 | 0.10 | 0.44 | 0.77 | 0 | 0 | `REPARATION_blast` |
| 1 | 68 | 1 | 30 | 0.03 | 0.99 | 0.97 | 0.50 | 0.50 | 0.06 | 0.69 | 0 | 0 | `Ribo-TISH` |
| 24 | 65 | 4 | 7 | 0.77 | 0.94 | 0.23 | 0.86 | 0.14 | 0.81 | 0.89 | 0 | 0 | `DeepRibo` |
| 5 | 58 | 11 | 26 | 0.16 | 0.84 | 0.84 | 0.31 | 0.69 | 0.21 | 0.63 | 0 | 0 | `IRSOM` |
| 9 | 48 | 21 | 22 | 0.29 | 0.70 | 0.71 | 0.30 | 0.70 | 0.30 | 0.57 | 0 | 0 | `SPECtre` |

##### D.5 Novel sORF detection

**Table S14:** Numbers of novel sORFs detected for all tools. The study of Weaver *et al* [2] could experimentally identify 33 novel sORFs. The novel ORFs detection power for each tool was evaluated by applying our benchmark pipeline on the *E. coli* Ribo-seq library described in [2]. This resulted in the here given numbers. Note: `DeepRibo` predicted in total 19, but using a cutoff 18 sORFs remain. Since no RNA library is available we could not test the novel ORF detection for `IRSOM`.

| Tool | `DeepRibo` | `REPARATION_blast` | `Ribo-TISH` | `SPECtre` |
|---|---|---|---|---|
| Novel sORFs | 18 | 2 | 0 | ? |

#### E Evaluation of key results

To follow the good practice benchmark suggestion proposed by Mangul *et al.* [6] we adapted one of their suggested example summary figures [7] for our benchmark scenario. Here, we propose a straightforward evaluation system to summarize the performance of all evaluated tools. The evaluation results are visualized in figure 5 of the main text. The evaluation system reveals the performance of each tool for each categories. We rate the tools as follows: a violet circle for superior performance, a light blue circle for satisfactory performance, and a dark blue circle for unsatisfactory performance. How each tool is rated for each category is described in the following.

##### E.1 Predictive power for the *translatome* set:

To evaluate the predictive power of each tool, we averaged the AUC values of the PRC of all *translatome* benchmark sets. `Ribo-TISH` and `IRSOM` achieved unsatisfactory results, with average AUCs of 0.69 and 0.70, respectively. `DeepRibo` and `REPARATION_blast` both achieved a superior predictive power, with AUCs of 0.94 and 0.91. The AUC values can be found in Table 3 of the main document.

Superior : AUC higher on average 0.90
Satisfactory : AUC higher on average 0.80
Unsatisfactory : AUC lower on average 0.80

##### E.2 Predictive power inside and outside of operons:

Averaging the AUCs for the detection of ORFs inside or outside of operons, respectively, led to the following results: `Ribo-TISH` 0.67 and 0.72, `DeepRibo` 0.93 and 0.95, `REPARATION_blast` 0.91 and 0.91, `IRSOM` 0.68 and 0.75

Superior : AUC higher on average 0.90
Satisfactory : AUC higher on average 0.80
Unsatisfactory : AUC lower on average 0.80

##### E.3 Prediction of novel sORFs:

The predictive power of finding novel ORFs was tested using 33 verified novel ORFs outside of the annotation[8], see main text. Only `DeepRibo` was able to find 18/19 of the 33 novel ORFs and has therefore a Satisfactory predictive power. `REPARATION_blast` with only two and `Ribo-TISH` and `IRSOM` with which found non of the novel ORFs show a unsatisfactory performance.

| | |
|---|---|
| Superior | : 20 novel ORFs detected |
| Satisfactory | : 10 novel ORFs detected |
| Unsatisfactory | : less than 10 novel ORFs detected |

##### E.4 Run time:

The run time comparison can be found in the main document (Table 7). We evaluated tools using single- and multithreading. Since not every tool is supports multithreading, we take the minimum run time for each tool of either single- or multi threading. Be aware that the original `REPARATION` tool should be faster than `REPARATION_blast` because of the use of `ublast` in `REPARATION_blast`.

| | |
|---|---|
| Superior | if test data was computed in less than 30 minutes |
| Satisfactory | if test data was computed in less than 2 hours |
| Unsatisfactory | if test data was computed in more than 2 hours |

##### E.5 Memory:

The memory comparison can be found in the main document (Table 7). We evaluated tools using single- and multithreading. Since not every tool supports multithreading, we take the minimum run time for each tool from either approach.

| | |
|---|---|
| Superior | : 2 or less GB |
| Satisfactory | : 4 or less GB |
| Unsatisfactory | : more than 4 GB |

##### E.6 Applicability:

The applicability describes how universally the tool can be applied. In the following we list several applicability criteria. If a tools fulfills the criteria it gets one point. Based on the amount of points each tool achieved the applicability is evaluated:

1. Can use replicates
2. Is deterministic
3. Outputs a standard file format (gff/bed/ ...)
4. Uses unit testing or some other correctness declaration
5. Stable results throughout different organisms

In the following, we describe how each tool was rated for applicability:

| | |
|---|---|
| Superior | : 4 or more points fulfilled |
| Satisfactory | : 2 or more points fulfilled |
| Unsatisfactory | : 1 or less points fulfilled |

**Table S15:** Applicability scoring table showing: (1) Can use replicates, (2) Is deterministic, (3) Outputs a standard file format, (4) Uses unit testing or some other correctness declaration, (5) Stable results throughout different organisms.

| tool | 1 | 2 | 3 | 4 | 5 | total |
|---|---|---|---|---|---|---|
| Ribo-TISH | 1 | 1 | 0 | 0 | 1 | 3 |
| DeepRibo | 0 | 1 | 1 | 0 | 1 | 3 |
| REPARATION_blast | 0 | 0 | 0 | 0 | 1 | 1 |
| IRSOM | 0 | 1 | 0 | 0 | 1 | 2 |
| SPECtre | 0 | 1 | 0 | 0 | 1 | 2 |

##### E.7 Usability:

User friendliness or usability is one of the key factors on how convenient it is for the users to apply the tool. In the following, we describe several usability criteria, each giving a single point if it is fulfilled by the tested tool.

1. Hosting software on a website with predicted long-term accessibility (e.g., GitHub)
2. Installation via package managers (e.g., Bioconda)
3. Provides an example dataset for testing
4. Version control (changelog)
5. Documentation: parameters
6. Documentation: input
7. Documentation: output
8. Documentation: dependencies
9. Open source

In the following, we describe the rating of the tools' usability:

**Table S16:** Usability scoring table showing: (1) Long-term accessible website, (2) Package managers, (3) Example data, (4) Version control, (5) Documentation: parameter, (6) Documentation: input, (7) Documentation: output, (8) Documentation: dependencies, (9) Open source. * *REPARATION* was using a proprietary, closed source sequence search tool that we replaced with the free and open source tool blast, to make the software usable without fee. This version is called REPARATION_blast.

| tool | 1 | 2 | 3 | 4 | 5 | 6 | 7 | 8 | 9 | total |
|---|---|---|---|---|---|---|---|---|---|---|
| Ribo-TISH | 1 | 1 | 0 | 1 | 1 | 1 | 1 | 1 | 1 | 8 |
| DeepRibo | 1 | 0 | 1 | 1 | 1 | 0 | 1 | 1 | 1 | 7 |
| REPARATION_blast | 1 | 1 | 1 | 0 | 1 | 1 | 0 | 1 | 1* | 7 |
| IRSOM | 1 | 0 | 1 | 0 | 0 | 1 | 0 | 1 | 1 | 5 |
| SPECtre | 1 | 0 | 1 | 1 | 1 | 1 | 1 | 1 | 1 | 8 |

| | |
|---|---|
| Superior | : 8 or more points fulfilled |
| Satisfactory | : 4 or more points fulfilled |
| Unsatisfactory | : 3 or less points fulfilled |
